# Comparative Transcriptome Profiling of High and Low oil yielding *Santalum album* L.

**DOI:** 10.1101/2021.05.12.443750

**Authors:** Tanzeem Fatima, Rangachari Krishnan, Ashutosh Srivastava, Vageeshbabu S. Hanur, M. Srinivasa Rao

**Affiliations:** Genetics and Tree Improvement Division, Institute of Wood Science and Technology Bangalore-India-560003; Laboratory for Structural Biology and Biocomputing Department of Computational and Data Sciences, Indian Institute of Science. Bangalore-India-560012; Department of Biotechnology, Indian Institute of Horticultural Research Hessarghatta, Bangalore 560089; Forest Development Corporation of Maharashtra Limited Nagpur-India-440036

**Keywords:** DEGs, Gene ontology, KEGG, qRT-PCR, *Santalum album*, Transcriptome analysis

## Abstract

East Indian Sandalwood (*Santalum album* L.) is highly valued for its heartwood and its oil. There have been no efforts to comparative study of high and low oil yielding genetically identical sandalwood trees grown in similar climatic condition. Thus we intend to study a genome wide transcriptome analysis to identify the corresponding genes involved in high oil biosynthesis in *S. album*. In this study, 15 years old *S. album* (*Sa*SHc and *SaS*Lc) genotypes were targeted for analysis to understand the contribution of genetic background on high oil biosynthesis in *S. album.* A total of 28,959187 and 25,598869 raw PE reads were generated by the Illumina sequencing. 2.12 million and 1.811 million coding sequences were obtained in respective accessions. Based on the GO terms, functional classification of the CDS 21262, & 18113 were assigned into 26 functional groups of three GO categories; (4,168; 3,641) for biological process (5,758;4,971) cellular component and (5,108;4,441) for molecular functions. Total 41,900 and 36,571 genes were functionally annotated and KEGG pathways of the DEGs resulted 213 metabolic pathways. In this, 14 pathways were involved in secondary metabolites biosynthesis pathway in *S. album.* Among 237 cytochrome families, nine groups of cytochromes were participated in high oil biosynthesis. 16,665 differentially expressed genes were commonly detected in both the accessions (*Sa*Hc and *Sa*SLc). The results showed that 784 genes were upregulated and 339 genes were downregulated in *Sa*Hc whilst 635 upregulated 299 downregulated in *Sa*SLc *S. album*. RNA-Seq results were further validated by quantitative RT-PCR. Maximum Blast hits were found to be against *Vitis vinifera*. From this study we have identified additional number of cytochrome family in *Sa*Hc. The accessibility of a RNA-Seq for high oil yielding sandalwood accessions will have broader associations for the conservation and selection of superior elite samples/populations for further genetic improvement program.

## Introduction

East Indian Sandalwood (*Santalum album* L; Family; Santalaceae) is evergreen hemi-parasitic perennial tree. *S. album* trees are found in semi-arid regions from India to the South pacific and the northern coast of Australia besides the Hawaii islands [1]. The economic value of sandalwood depends on the quantity of heartwood and its essential oil extracted from the heartwood as well roots of the mature trees of santalum spps. [2,3,4,5]. It has been used for perfumery, cosmetics, pharmaceutical, religious and cultural purposes over centuries [6]. Indian government categorized *S. album* as one of 32 recognized medicinal plant (Gowda, 2011) [7]. The essential oil is very important trait, which is subjected to host species, soil type, climate effects and elite germplasm [8,9,10,11,12]. However the limited oil yield of sandalwood restricts the demand of oil. The sandalwood oil formation is independent of heartwood growth and it was assumed that constant amount of oil being formed nevertheless of trees/heartwood growth, similar age of trees and with the smaller diameter heartwood consisting trees may tend to have greater percentage of oil. The quality of oil is largely defined by the percentage of different fragrant sesquiterpenes within the oil, especially α and β santalol [5]. Out of other santalum species, *S. album* is valued as a source of high content of oil as it has high level of α and β santalol and it shows low variability in oil composition across its natural range [13]. Due to international demand for sandalwood heartwood and its oil, over the recent times *S. album* has been considered as private investment to develop a sandalwood industry [14]. Excessive harvest, habitat destruction and lack of pest management system, global sandalwood resources are threated globally which indicated the large-scale shortage and escalation the market price of sandalwood products [15,4,16]. Realizing the sharp decline in the sandalwood population, the Karnataka and Tamil Nadu Forest department amended the sandalwood act in 2001 and 2002 and declared the private sandalwood growers himself an owner of the sandalwood as per the amended Act. Further, Govt. of Karnataka made an amendment on the sale of sandalwood through Forest department and Government, Departments to eliminate the clandestine trade and to encourage farmers to take cultivation of Sandalwood on commercial scale during the last few years [7]. Due to the amendment, many of the private organizations and farmers have started raising sandalwood cultivation on their private/farm lands. Since sandalwood plantation is long term high investment by the farmers and forest department, so it is essential to identify and supply superior quality planting material to optimize the high economic returns than their investment.

The breeding improvement is little due to its long generation time and lack of information about high oil yielding accessions/populations. Considering the constant increasing the global demand for sandalwood oil and genetic improvement purposes, the identification of factors regulating these qualitative and quantitative variations in oil is a critical issue. It was hypothesized that accumulation of sandalwood oil is a complex and dynamic process, which influenced by multiple genetic and environmental factors (17). Candidate oil biosynthesizing genes, multiomics, trait associated mapping have been performed to investigate the mechanism of oil biosynthesis and accumulation. With the advancement of high throughput sequencing technology, several transcriptome profiling of studies have been carried out in sandalwood [18,19,20,21]. Although earlier studies showed that sandalwood oil biosynthesis pathways, identification of key oil biosynthesis genes (Cytochrome P450, Sesquisabinene synthases, and Sesquiterpene synthases), there are very few references available on transcriptomic oil biosynthesis regulation and accumulation. As such there is no any studies pertaining on transcriptomic regulation of sandalwood clones grown in identical environmental conditions. In this study, we performed comparative transcriptomic profiling of two identical accessions that differ significantly in oil content to understand the dynamic regulation of high and low oil accumulation. Understanding the high and low oil variants of the trees, as even a slight percentage improvement in sandalwood oil content will lead to significant value [22,23]. Our results provide new insight for better understanding of how to achieve more sandalwood oil production by manipulation of core pathways and gene involved.

## Materials and Methods

### Sampling site

The selection of *S. album* samples for transcriptome analysis was grounded on three factors (1) known age and (2) grown in identical environmental condition (3) diseased free trees. Therefore we selected 15 year old *S. album* trees grown in Institute of Wood Science and Technology (13.011160°N 77.570185°E) Bangalore Karnataka and collected samples in the month of August 8^th^ 2018.

### Sample collection

For oil estimation and RNA isolation, the wood samples were collected up to GBH at 1.37 M by using conventional drilling increment borer (leaf materials were takes as a positive control in RNA extraction process). The core samples were marked as transition zone, heartwood and sapwood and frozen into liquid nitrogen. The samples were immediately stored in dry ice box and shipped to the Eurofins laboratory. Before RNA extraction from the core samples, the oil quantity and quality was estimated by UV-spectrophotometer followed by GC-MS analysis. Based on the oil variability in terms of high and low oil-yielding (*Sa*SHc and *Sa*SLc) samples were selected for *De novo* transcriptome analysis S1 Table.

### RNA isolation, cDNA library preparation and Sequencing

The total RNA was extracted from transition zones of the selected cores and leaf (+ control) samples by using modified CTAB and LiCl method [24,25] The quality of isolated RNA measured by UV spectrophotometer at 260/280and 260/230 nm wavelengths and 1% agarose gel electrophoresis followed by measuring RNA concentration using a 2100 Bioanalyzer (Agilent Technologies). The concentration of RNA was obtained in *Sa*SHc 1460.90 ng/*μ*l and in *Sa*SLc 12.65 ng/μl. The mRNA from the total RNA was extracted by using the poly-T attached magnetic beads, followed by fragmentation process. The cDNA library of *S. album* was constructed using 2 µL of total purified mRNA from each sample by using Illumina TruSeq stranded mRNA preparation kit. 1st strand cDNA conversion was carried out by using Superscript II and Act-D mix to facilitate RNA dependent synthesis and then second strand was synthesized by using second strand mix. The dscDNA was purified by using AMPure XP beads followed by A-tailing adapter ligation. The libraries were analyzed through 4200 TapeStation system (Agilent Technologies) by using high sensitivity D1000 screen tape. The Pairing end Illumina libraries were loaded on NextSeq500 for cluster generation and sequencing. Total two RNA libraries were generated with the Paired end sequencing. To obtain high quality concordant reads the sequenced raw data were processed by Trimmomatic v0.38 [26]. In-house script (in python and R) software was used to remove adapters, ambiguous reads and low quality sequences and the high quality paired-end reads were used for *De novo* Transcriptome assembly. RNA-Seq data were produced in FASTQ format and the whole sequence reads archive (SRA) database has been deposited in NCBI under Biosample accession: SAMN1569426 SRA accession number: PRJNA648820.

### *De Novo* Transcriptome Assembly, Unigenes classification and Functional Annotation

Trinity *de novo* assembler (v2.5) [27] was used to assemble transcripts from pooled reads of the samples with a kmer_25 and minimum contig length value up to 200 bp. The assembled transcripts were then further clustered into unigenes covering >90% at the 5X reads by using CD-HIT-EST-4.5.4 software [28] for further downstream analysis. Coding sequences (open reading frames, ORFs) within the unigenes (default parameters, minimum of 100 amino acid sequence) were predicted by TransDecoder v5.0. The longest ORFs were then subjected to BLAST analysis against PSD, UniProt, SwissProt, TrEMBL, RefSeq, GenPept and PDB databases to obtain protein information resource (PIR) for the prediction of coding sequences by Blast2GO software program [29].

### Functional Annotation

The functional annotation of genes was performed by DIAMOND (BLASTX compatible aligner) program software [30]. The functional identification of coding sequences in biological pathways of the respective sample reads was assigned to reference pathways in KEGG (Eukaryotic database). The output of KEGG analysis included KEGG orthology, corresponding enzyme commission (EC) numbers and metabolic pathways of predicted CDS by using KEGG automated annotation server KAAS (http://www.genome.jp/kaas-bin/kaas_main) [31].

### Differential gene expression analysis

The differential expressed genes (DEGs) were identified between the corresponding samples by implementing a negative binomial distribution model in DESeq package (v.1.22.1_http://www.huber.embl.de/users/anders/DESq) [32]. The combination for differential analysis was calculated as *Sa*SH1 (high oil yielding) vs *Sa*SL1 (low oil yielding) *S. album.* To analyze the differentially expressed genes, two software’s (heatmap, and Scatter plot) were used to predict upregulated and downregulated genes in *S. album.* A heat map was constructed by using the log-transformed and normalized value of genes based on Pearson uncentered distance and average linkage method. The most similar transcriptome profile calculated by a single linkage method, a heatmap were generated, correlating sample expression profiles into colors. The heatmap shows the level of gene expression and represented as log2 ratio of gene abundance between high and low oil yielding samples. An average linkage hierarchical cluster analysis was performed on top 50 differentially expressed genes using multiple experiments viewer (MeV v4.9.0) [33]. The color represents the logarithmic intensity of the expressed genes. Relatively high expression values were showed in red (identical profiles) and low expression values were showed in green (the most different profiles). The scatter plot is used for representing the expression of genes in two distinct conditions of each sample combination i.e., high and low oil yielding clones. It helps to identify genes that are differentially expressed in one sample with respect to the corresponding samples. This allows the comparison of two values associated with genes. The vertical position of each gene in form the of dots represents its expression level in the high oil yielding samples while the horizontal position represents its expression level in the treated samples. Thus, genes that fall above the diagonal are over-expressed and gene that fall below the diagonal are under expressed as compared to their median expression level in experimental grouping of the experiment.

### Quantitative RT-PCR Analysis

Quantitative Real Time (qRT) PCR was performed by using SYBR Green PCR master mix kit in a stepOnePlus Real Time PCR system (Applied Biosystem by Life Technologies, USA). To validate the gene expression profiles identified by RNA-Seq. 2 μg of RNA was reverse transcribed in a 20 μL volume with RT PCR master mix (TaKaRa) as per the manual instruction. Six gene (*Sa*MTPS, *Sa*FPPS, *Sa*DSX, *Sa*GGPS, *Sa*GPS, and SaCYP450) specific primers were predicted using by the online tool Primer3 version 0.4.0 and synthesised at (Eurofins India Pvt. Ltd). The sequence of primers with a melting temperature between 60-61 ^0^C and a PCR product range of 151-229 bp were listed in S2 Table. Actin was used as a reference gene. qRT-PCR was performed with step One Real time PCR system (Applied Biosystems, Thermofisher Scientific). The qRT-PCR reaction systems were as follows: 95^0^ C for 20 s, followed by 40 cycles of 95^0^C for 5 s, 60^0^C for 30s and 72^0^ C 40 sec. The fluorescence data were collected and analysed with Step One analysis software.

## Results

### Qualitative Analysis of *S. album* oil

The selected core samples were quantitatively and qualitatively analyzed. The total oil percentage was found 4.96% and 0.93% for respective samples. Along with the oil content, α/β-santalol variation in *Sa*SHc 59.30/32.21 and in *Sa*SL 49.52/26.60 was observed S1 Table.

### Library construction and Transcriptome Sequencing

A total of 38,785326 (*Sa*SH) and 35,94,4784 (*Sa*SL) raw PE reads were generated from the Illumina sequencing of *S. album* Table 1. After removing adapters containing >5% unknown nucleotide sequences, ambiguous reads and low quality reads (reads with more than 10% quality threshold (QV) <20phred score) 28,959187 and 25,598869 were obtained to respective samples. The total clean bases for *Sa*SHc were 4.4 GB with 47.67% GC and 3.8 GB with 48.62% GC content for *Sa*SLc. 141,781 clean pair-end reads were assembled into pooled non-redundant putative transcripts with the mean length of 1,149 bp followed by N50: 2,044. The obtained transcript length ranged from 201 to 15,872 S3 Table. The transcripts were assembled into 31,918 unigenes with the mean length and N50 length 1,739 2,272 respectively S3 Table. Of the unigenes we found 11.85% (3,785) 200-500 bp in length, 19.06% (6,085) were 500-1000 bp in length, 36.28% (11,582) were 1000-2000 bp in length, 19.35% (6179) 2000-3000 bp in length, 8.42% (2688) 3000-4000 bp in length, 2.96% (946) 4000-5000 bp in length and 2.04% (653) exceeded 5000 bp (Table S3). A total number of coding sequences (CDS) in pooled samples were found 2.271 million with total 2.810 billion bp. S3 Table. Sample wise number of CDS was in *Sa*SHc and *Sa*SLc was 2.12 million and 1.811 million followed by total CDS base length 2.657 billion in *Sa*SHc and 2.307 billion S3 Table.

**Table 1.** Summary of cDNA library, RNA-Seq and *de novo* sequence assembly of combined (*Sa*SHc and *Sa*SLc) *S. album*

| **Description** | ***SaSHc*** | ***SaSLc*** |
| --- | --- | --- |
| cDNA library size (bp) | 252-662 | 232-571 |
| Average cDNA size (bp) | 416 | 375 |
| No. of PE reads | 2.99 billion | 2.55 billion |
| Number of bases | 435.67 billion | 384.98 billion |
| Total data in GB | 4.4 | 3.89 |

### Gene Functional Annotation and Classification

Total 22,710 CDS were BLAST and 20,842 CDS were annotated by NCBI databases (Table S3). In case of *Sa*SHc and *Sa*SLc 20,262 and 18,113 genes were studied for Gene Ontology (GO). Based on the transcripts distribution, the assembled CDS were assigned into 26 functional groups of three GO categories: (i) Biological process (*Sa*SHc 4,168; *Sa*SLc 3,641) (ii) Molecular functions (*Sa*SHc 5108; *Sa*SLc 4,441) and (iii) Cellular components (*Sa*SHc 15,758; *Sa*SLc 4,971) (Table 2) (Fig 1. A, B, C). GO annotations for molecular functions (*Sa*SHc 13; *Sa*SLc 12), biological process (*Sa*SHc; 21, *Sa*SLc; 22) and cellular component analysis *Sa*SHc (16) and *Sa*SLc (17) were plotted by WEGO plotting tool. These domains were further containing Cellular component and in Molecular functions followed by Biological process respectively. The number of differential expressed genes (DEGs) in biological regulation terms was observed 5,108 in *Sa*SHc and 4,442 in *Sa*SLc. Data showed that prominent GO terms in biological process were metabolic process, cellular process, biological regulation, localization, stimulus, cellular component organization or biogenesis and signaling. Similar result was observed in cellular components *viz, Sa*SHc (4,168) and *Sa*SLc (3,642). In cellular components, majority of GO terms was related to cell, cell part organelle, membrane enclosed lumen, membrane and protein containing complex related genes was overrepresented in *Sa*SHc. In molecular function, the number of DEGs were involved in GO terms was 5,758 in *Sa*SHc and 4,972 in *Sa*SLc. The DEGs were prominently participated in catalytic activity, binding, transport activity, molecule carrier activity, antioxidant activity, and signal transducer activity. Among cellular components, cytosol, intracellular part, cytoplasmic fraction and cytoplasm were overrepresented in *Sa*SHc as compared to *Sa*SLc accessions. High number of genes was found in *Sa*SHc (41,900 genes) compared to *Sa*SLc (36,571 genes) that was further classified into biological process, cellular component and molecular functions. Highest number of genes was functionally annotated and was observed in biological process (*Sa*SH 16,361) and (*Sa*SL 14,459) followed by molecular function (Fig 2 A & B).

**Figure 1 (A).**
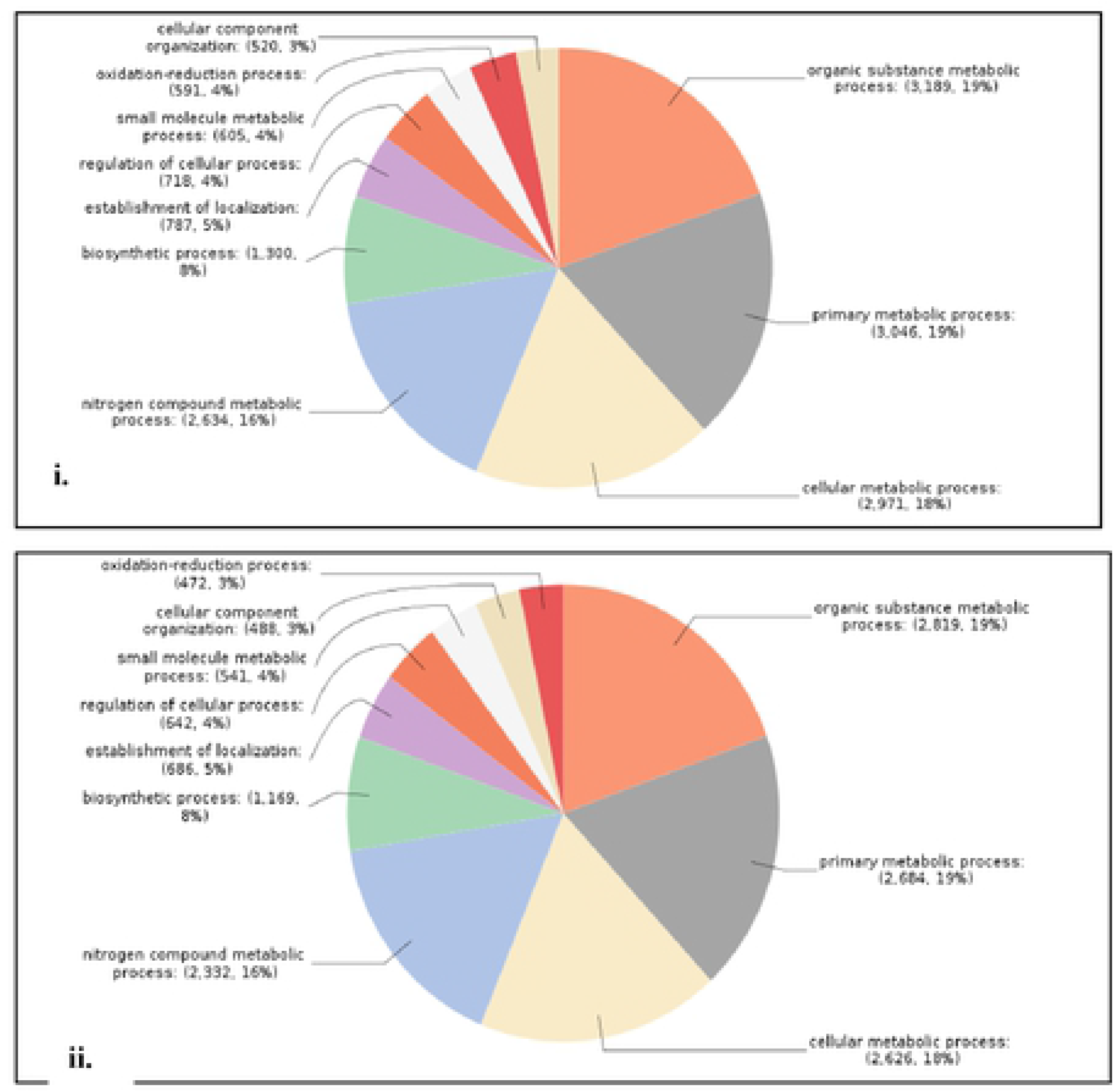
Comparative Go biological regulation **i.** High oil yielding (*Sa*SHc) and **ii.** low oil yielding (*Sa*SHc) in *S. album.*

**Figure 1 (B).**
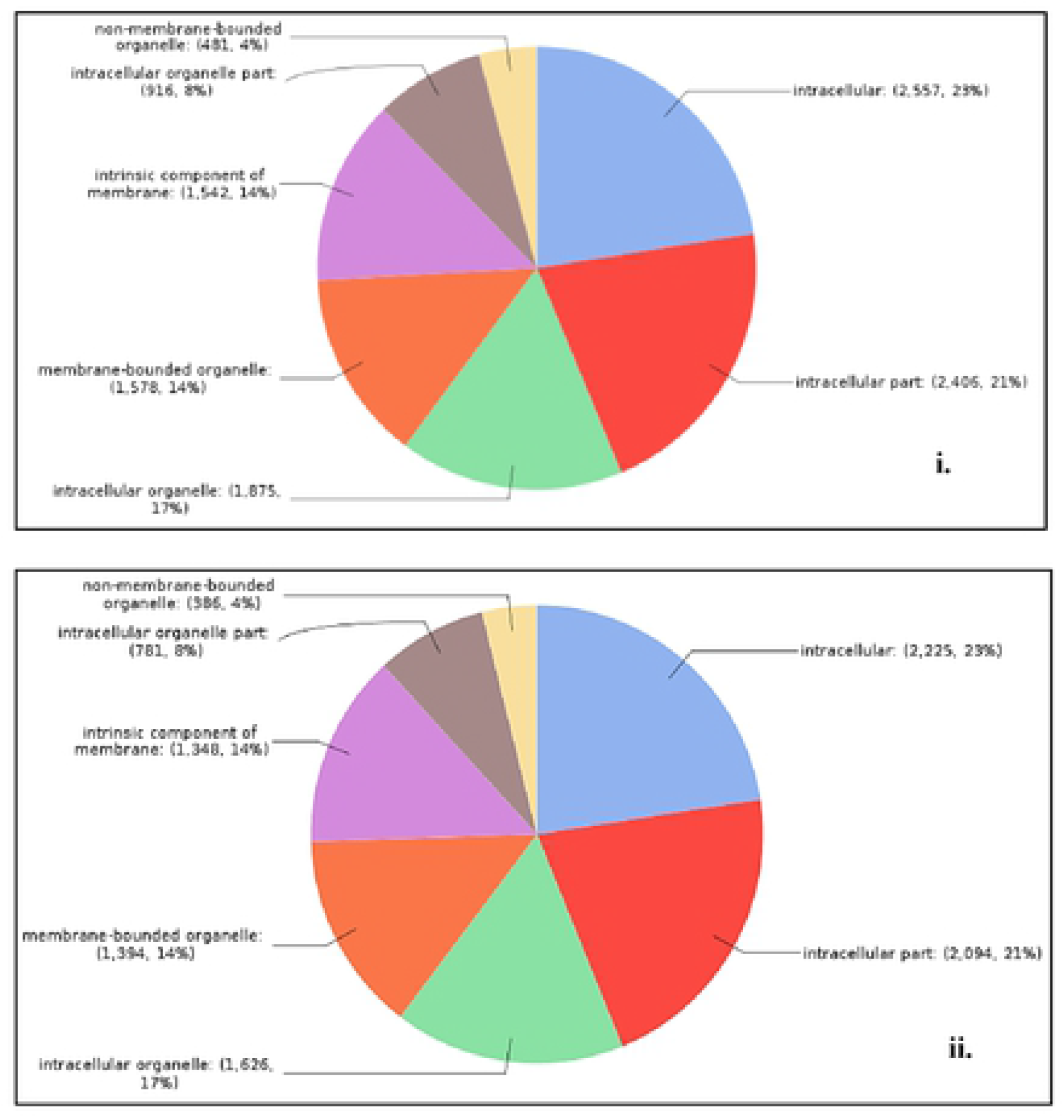
Comparative Go Cellular component **i.** High oil yielding (*Sa*SHc) and **ii.** low oil yielding (*Sa*SHc) in *S. album.*

**Figure 1 (C).**
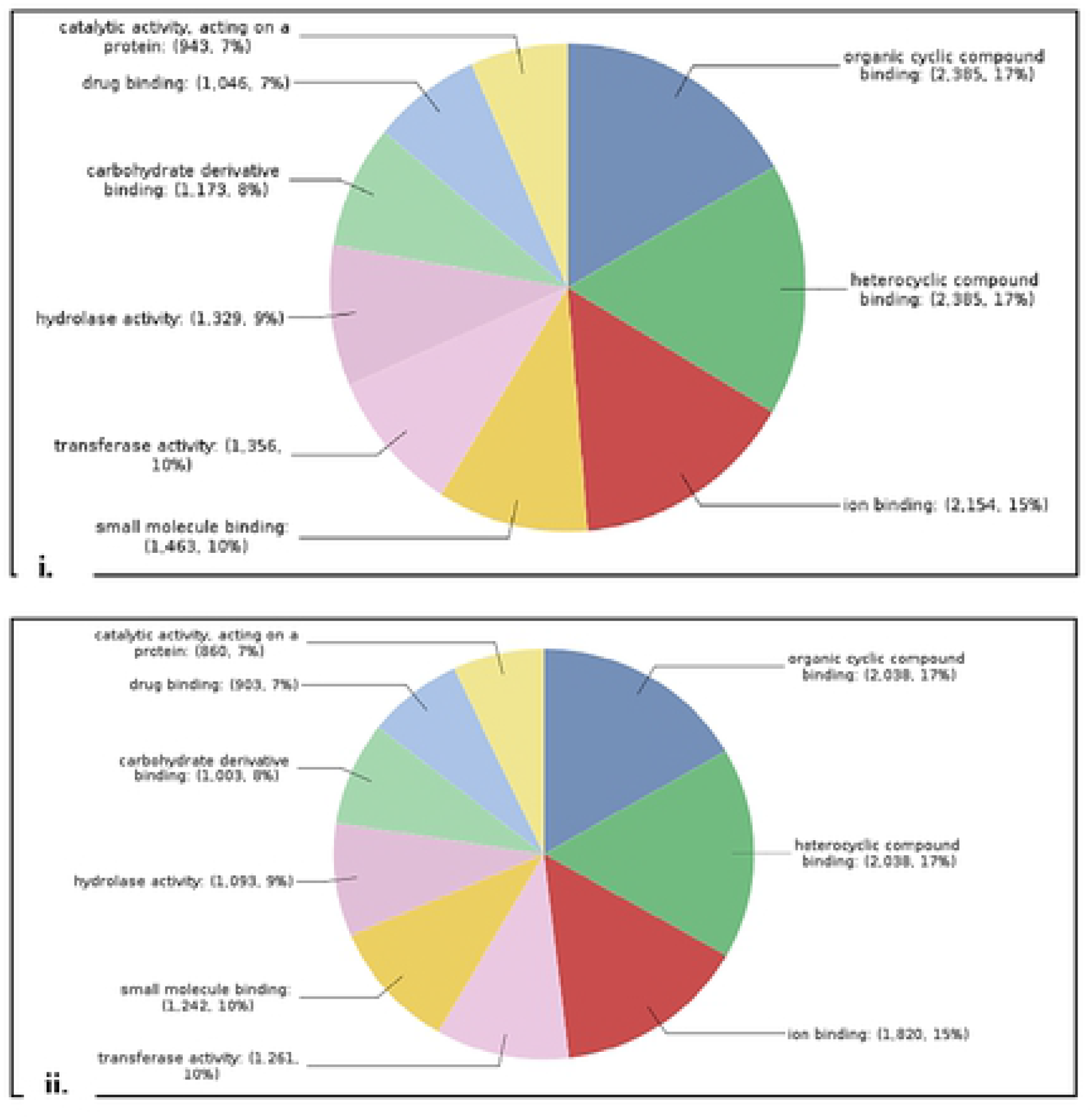
Comparative Go Molecular function between **i.** High oil yielding (*Sa*SHc) and **ii.** low oil yielding (*Sa*SLc) in *S. album*.

**Figure 2 (A).**
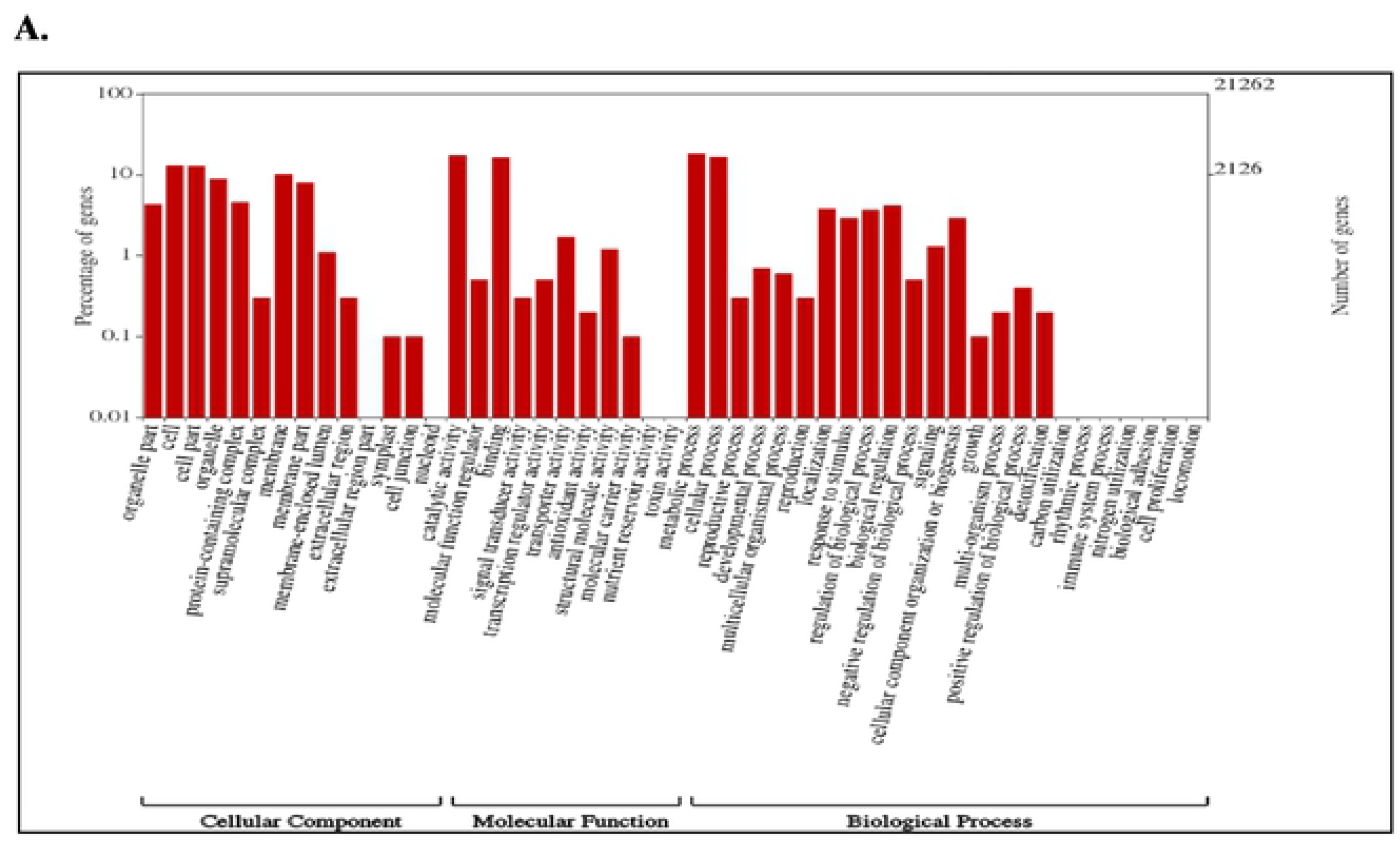
Histogram of gene ontology classification (Wego plot); High oil yielding Sandalwood (*Sa*SHc).

**Figure 2 (B).**
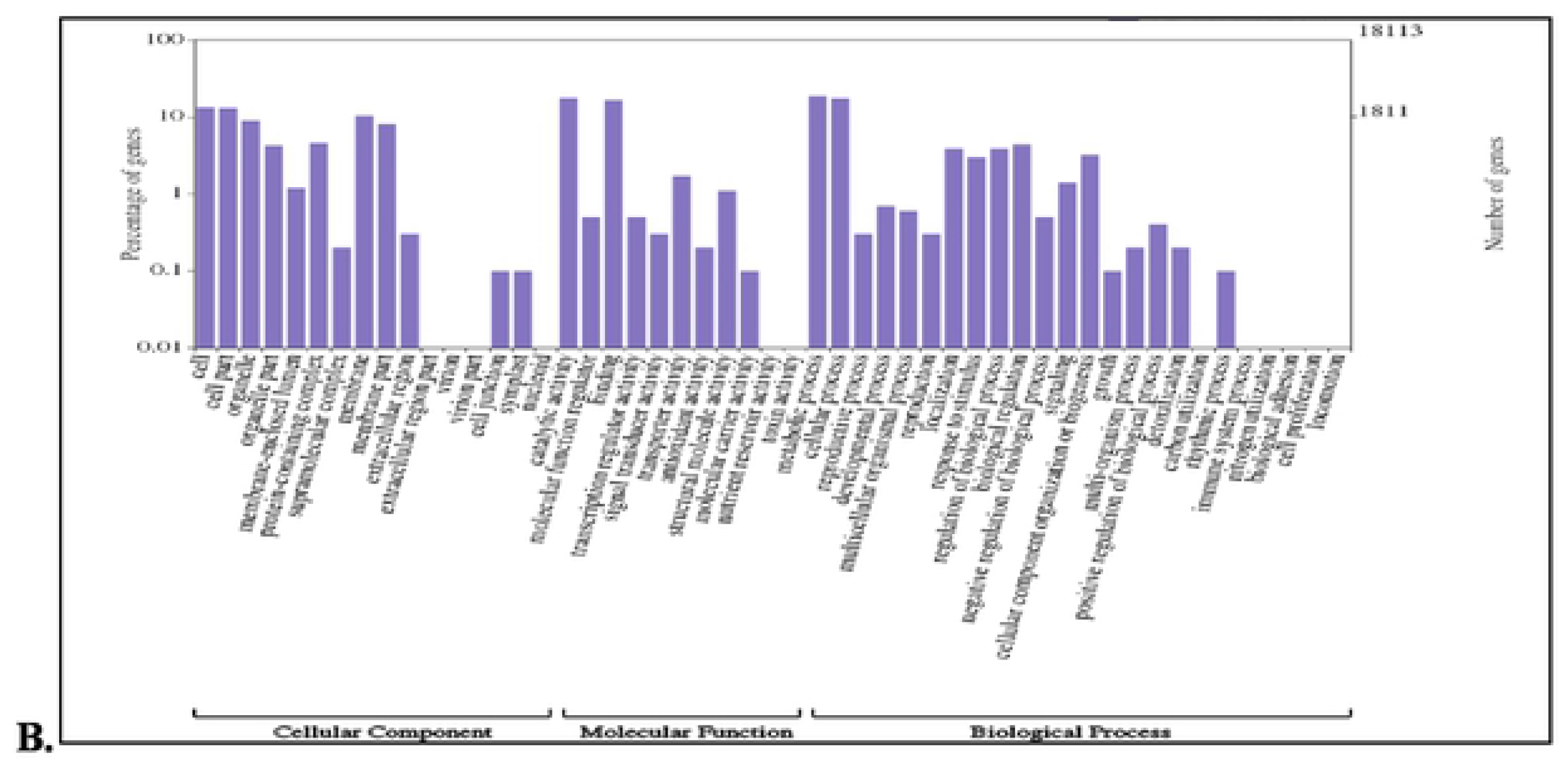
Histogram of gene ontology classification (Wego plot); Low oil yielding Sandalwood (*Sa*SLc).

**Table 2.** Samples wise Gene ontology (GO) category distribution of coding sequences (CDS) in *S. album*

| **SI No.** | **Biological Process** | **Cellular Component** | **Molecular Function** |
| --- | --- | --- | --- |
| **SH1** | 5,108 | 4,168 | 5,758 |
| **SL1** | 4,441 | 3,641 | 4,971 |

### Kyoto Encyclopedia of Genes and Genomes (KEGG) pathway mapping

Significant DEGs between *Sa*SHc and *Sa*SLc were mapped to reference canonical pathways in KEGG database. A total of 6,159 and 5,554 CDS of *Sa*SH and *Sa*SL were found to be categorized into 24 major KEGG pathways and were grouped in five main categories (Table 3). All assembled unigenes were subjected to further functional prediction and classification by KEGG Orthology (KO) database. Results showed 6,159 and 5,554 unigenes involvement in 24 groups in the KO database in respective samples and further subcategorized into 213 metabolic pathways (Table 3; Fig S1 A, B; S2 & S3). KEGG metabolite pathways represented 10 major pathways like metabolism, terpenoid synthesis, amino acid metabolism, purine metabolism, pyrimidine, transcription, translation, amino acyl-tRNA biosynthesis, DNA replication and membrane transport in sandalwood (Table 4). The EC numbers were classified in KEGG pathways, enabling the presentation of enzymatic functions in the context of the metabolic pathways. Among the identified pathways, secondary metabolite-flavonoid, and terpenoid related transcripts were over-represented (Table 4).

**Table 3.** Comparative KEGG pathway classification of coding sequences in high oil (SH1) and low oil (SL1) yielding *S. album*

| Pathways | SaSHc | SaSLc |
| --- | --- | --- |
| **Metabolism** |  |  |
| Carbohydrate Metabolism | 556 | 494 |
| Energy metabolism | 323 | 281 |
| Lipid metabolism | 272 | 231 |
| Nucleotide metabolism | 162 | 147 |
| Amino acid metabolism | 393 | 362 |
| Metabolism of other amino acids | 156 | 138 |
| Glycan biosynthesis and metabolism | 99 | 88 |
| Metabolism of cofactors and vitamins | 218 | 193 |
| Metabolism of terpenoids and polyketides | 99 | 80 |
| Biosynthesis of other secondary metabolites | 86 | 77 |
| Xenobiotics biodegradation and metabolism | 85 | 57 |

**Table 3.** Comparative KEGG pathway classification of coding sequences in high oil (*Sa*SHc) and low oil (*Sa*SLc) yielding *S. album*

| Pathways | *SaSHc* | *SaSLc* |
| --- | --- | --- |
| **Environmental Information Processing** |  |  |
| Membrane transport | 34 | 30 |
| Signal transduction | 597 | 645 |
| Signaling molecules and interaction | 0 | 1 |
| **Cellular Processes** |  |  |
| Transport and catabolism | 458 | 426 |
| Cell growth and death | 329 | 297 |
| Cellular community – eukaryotes | 94 | 87 |
| Cellular community – prokaryotes | 72 | 67 |
| Cell motility | 51 | 44 |
| **Genetic information** |  |  |
| Transcription | 321 | 301 |
| Translation | 739 | 652 |
| Folding, sorting and degradation | 551 | 526 |
| Replication and repair | 151 | 126 |
| **Organismal system** |  |  |
| Environmental adaptation | 264 | 253 |

**Table 4.** Sandalwood transcripts mapped to KEGG pathway (Top 10)

| SI No | KEGG pathways | SaSHc | SaSLc |
| --- | --- | --- | --- |
| 1. | Metabolism | 2981 | 2633 |
| 2. | Terpenoid synthesis | 216 | 181 |
| 3. | Amino acid metabolism | 557 | 495 |
| 4. | Purine metabolism | 130 | 119 |
| 5. | Pyrimidine metabolism | 82 | 73 |
| 6. | Transcription | 321 | 301 |
| 7. | Translation | 303 | 234 |
| 8. | Amino acyl tRNA biosynthesis | 46 | 42 |
| 9. | DNA replication | 27 | 26 |
| 10. | Membrane transport | 34 | 30 |

### DEGs involved in sandalwood oil biosynthesis in *S. album*

DEGs were further annotated with KEGG database to deep insight the gene products for metabolism and functions related genes in different classified pathways. We performed an enrichment analysis of gene ontology (GO) terms with high significance in the upregulated DEGs. To identify metabolic pathways, *Sa*SHc (297) and *Sa*SLc (259) DEGs were mapped. As a result, 14 major pathways have been shown to play important role in sandalwood oil biosynthesis. Most pathways were resulted to secondary metabolites biosynthesis and metabolism by cytochrome P450. In order to identify secondary metabolite biosynthesis pathways in sandalwood, 4,697 transcripts for *Sa*SHc and 4,134 for *Sa*SLc were plotted. In Terpenoid backbone biosynthesis (35;33), Monoterpenoid biosynthesis (2;1), Sesquiterpenoid and Tri-terpenoid biosynthesis (4;3), Diterpenoid biosynthesis (10;10), Polyprenoid biosynthesis (31;30), Flavone and Flavanol biosynthesis (3;2), Isoquinolene alkaloid biosynthesis (9;6), Stilbenoid diaryl-heptanoid and Gingerol biosynthesis (3;4), Tropane piperidine and pyridine alkaloid biosynthesis (11;18) and Carotenoid biosynthesis (21;15) genes were involved in *Sa*SHc and *Sa*SLc sandalwood accessions. Predominantly genes were involved in metabolism of xenobiotics by Cytochrome P450 (*Sa*SHc 34; *Sa*SLc 23) and leads to up-regulation metabolic pathways. All these Go terms can be connected with sandalwood oil biosynthesis through an enhanced production of gene products in *S. album* oil biosynthesis pathway (Table 5).

**Table 5.** Comparative analysis of DEGs involved in secondary metabolite biosynthesis pathway analysis of Kos in high oil (*Sa*SHc) and low oil (*Sa*SLc) yielding *S. album*

| Pathway | Kos |  | Pathway ID |
| --- | --- | --- | --- |
|  | *SaSHc* | *SaSLc* |  |
| Terpenoid backbone biosynthesis | 35 | 33 | Ko00900 |
| Monoterpenoid biosynthesis | 2 | 1 | Ko00902 |
| Sesquiterpenoid and triterpenoid biosynthesis | 4 | 3 | Ko00909 |
| Diterpenoid biosynthesis | 10 | 10 | Ko00904 |
| Polyprenoid biosynthesis | 31 | 30 | Ko00940 |
| Flavone and flavanol biosynthesis | 3 | 2 | Ko00944 |
| Isoquinolene alkaloid biosynthesis | 9 | 6 | Ko00950 |
| Drug metabolism: Cytochrome | 31 | 23 | Ko00982 |
| Metabolism of xenobiotics by Cytochrome P450 | 34 | 23 | Ko00980 |
| Stilbenoid diarylheptanoid and gingerol biosynthesis | 3 | 4 | Ko00945 |
| Tropane piperidine and pyridine alkaloid biosynthesis | 11 | 8 | Ko00960 |
| Carotenoid biosynthesis | 21 | 15 | Ko00906 |
| **Total** | **194** | **158** |  |

### Profiling of Differential Expressed Genes (DEGs) participated in sandalwood oil biosynthesis regulation

All stages of sandalwood oil biosynthesis were examined, and a comparative analysis was done using aligned reads and the transcripts were grouped based on their degree of expression (log2 FC). 16,665 differentially expressed genes were commonly detected in both the accessions (*Sa*Hc and *Sa*SLc). The results showed that 784 genes were upregulated and 339 genes were downregulated in high oil yielding accessions whilst 635 upregulated 299 downregulated in low oil yielding *S. album* accessions (Fig. 3). Gene expression pattern represented by Scatter plot showed a significant log 2FC>16.0; P value <0.005 for upregulated genes and log 2FC<0.40; P value <0.005 downregulated in case of *Sa*SHc sample. 4.39% genes were found upregulated and 1.87% was downregulated in total differentially expressed genes. The normalized gene expression values from both the samples were used to estimate a Euclidian distance matrix based on transcript describing the similarities between the *Sa*SHc and *Sa*SLc samples. Red dots represented the upregulated genes and green dots represented the down regulated in DGE combination Fig 4. Similar to scatter plot, based on their degree of expression (log2 FC) values heatmap were also used to generate DEGs pattern. Heatmap showed transcript abundance level and indicated a similarity gradient between the *Sa*SHc and *Sa*SLc accessions. In heatmap, gene expression was calculated in accordance with the method of FPKM, which takes into account the influence of both the sequencing depth and gene length on read count. In the FPKM distribution for selected samples, *Sa*SHc showed the highest probability density distribution of gene expression, whereas, *Sa*SLc displayed the lowest Fig 5. The transcripts, which were highly expressed, were annotated for each gene as a high number of fold change and measure primarily the relative change of expression level. The top 50 highly upregulated genes (log2 FC 9.285-4.65) were shown in heatmap (Fig 5). The transcriptional mining identified ten unigenes participated in sandalwood oil biosynthesis with the upregulated relative gene expression log2 FC *viz,* **(i)** Geranyl geranyl diphosphate synthase (GPS) (FC; 3.54), **(ii)** Geranyl diphosphate synthase (GGPS) (2.6), **(iii)** 3-hydroxy-3-methylglutaryl-coenzyme A reductase (HMG-CoA) (1.32), **(iv)** 1-Deoxy-D-xylulose-5-phosphate synthase (DXS) (0.675), **(v)** E-E, Farnesyl pyrophosphate synthase (E-E-FDS) (3.21), **(vi)** cytochrome P450 synthase (CYP450) (2.43) **(vii)** Farnesyl pyrophosphate synthase (FPPS) (1.86), **(viii)** Phenylalanine ammonia lyase (2.1) **(ix)** Monoterpene synthase (MTPS) (2.76), **(x)** 5-enolpyruvylshikimate 3-phosphate synthase (ESPS) (1.4) (Table 6; Table S4). Transcripts encoding *Sa*FPPS gene in *Sa*SHc showed 10 fold higher than *Sa*SLc accessions (Table 6; S4 Table).

**Figure 3.**
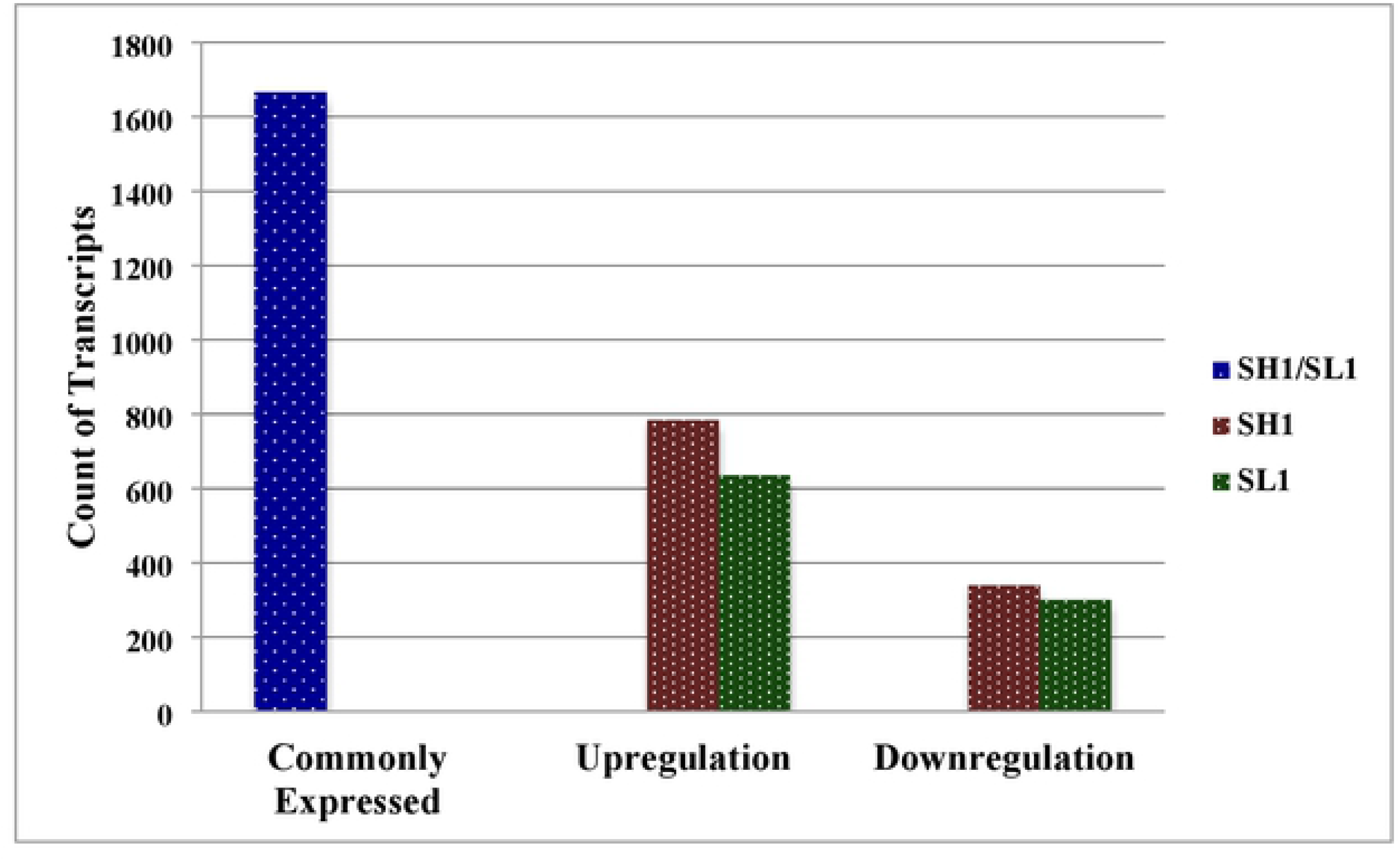
Identification of differentially expressed genes (DEGs) between *Sa*SHc and *Sa*SLc. Green Bar indicates commonly expressed DEGs. Blue and red bars represent upregulated and downregulated DEGs.

**Figure 4.**
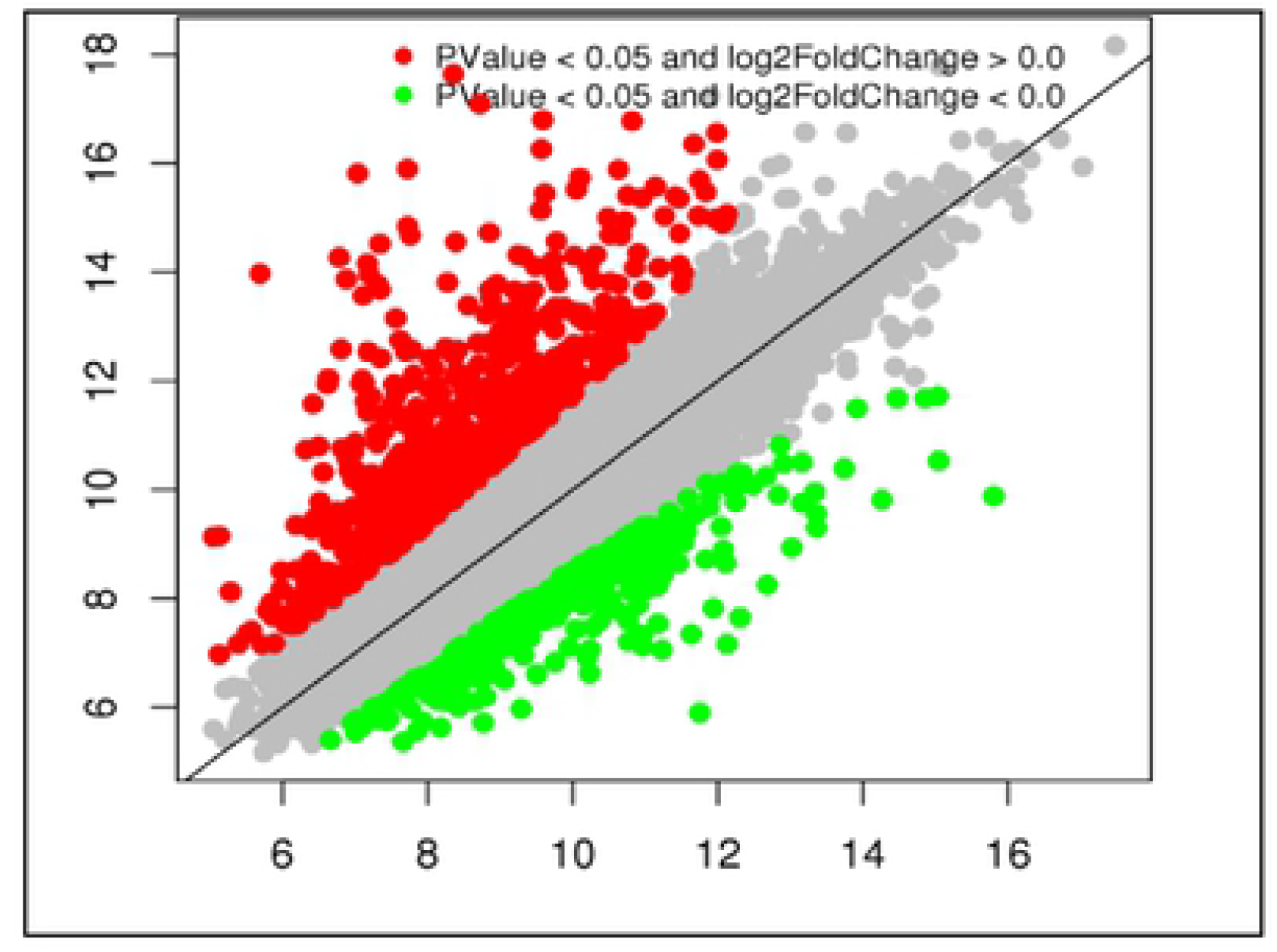
Visualization of differentially expressed gene transcription by Scatter plot between *Sa*SHc and *Sa*SLc samples.

**Figure 5.**
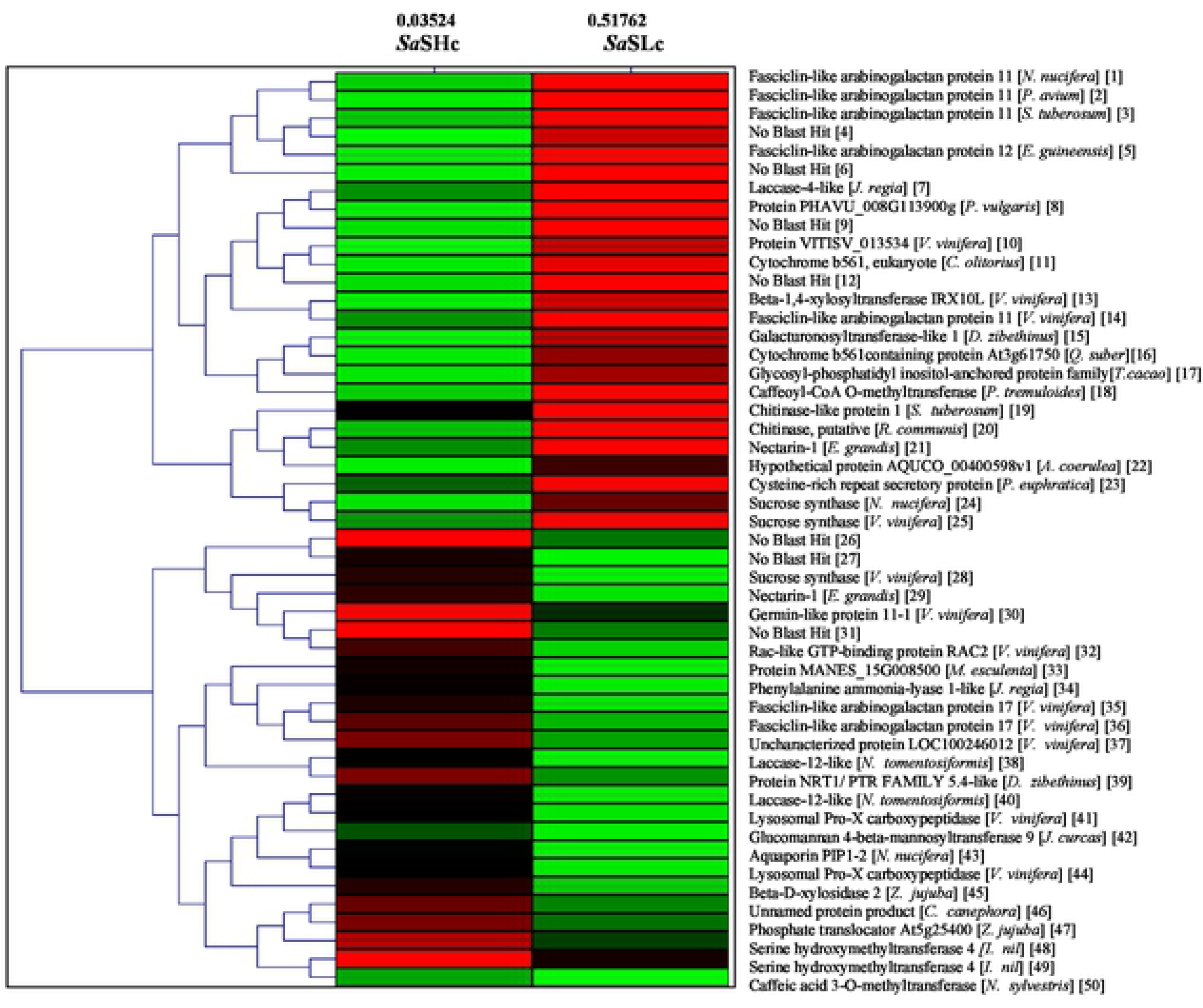
Heat map depicting the top 50 differentially expressed genes (significant); base Mean *Sa*SHc represents the normalized expression values for *Sa*SHc sample and base Mean and *Sa*SLc represents the normalized.

**Table 6.** Relative expression of high oil yielding (*Sa*SHc) genes, coding sequences, unigenes, transcripts, log2 fold change and regulation

| SI No. | Genic SSR primers | CDS | Unigenes | Transcripts | Log2fold change | Regulation |
| --- | --- | --- | --- | --- | --- | --- |
| 1. | Geranyl pyrophosphate synthase (GPS) | 2915 | 3614 | 38599 | 2.61 | Upregulation |
| 2. | Geranyl geranyl pyrophosphate synthase (GGPS) | 9544 | 11617 | 66429 | 3.54 | Upregulation |
| 3. | 3-Hydroxy-3-methylglutaryl-CoA reductase (HMG-CoA) | 11763 | 14481 | 75932 | 1.32 | Upregulation |
| 4. | 1-Deoxy-D-xylulose5-phosphate synthase (DXS) | 21435 | 27658 | 119568 | 0.67 | Upregulation |
| 5. | E, E, Farnesyl diphosphate synthase (E-E-FDS) | 7514 | 29031 | 123929 | 2.65 | Upregulation |
| 6. | Cytochrome P450 synthase (CYP450) | 6012 | 7205 | 5126 | 2.43 | Upregulation |
| 7. | Farnesyl pyrophosphate synthase (FPPS) | 1770 | 2262 | 32293 | 1.86 | Upregulation |
| 8. | Phenylalanine ammonia lyase (PAL) | 21850 | 28225 | 121398 | 3.55 | Upregulation |
| 9. | Monoterpene synthase (MTPS) | 1948 | 2474 | 33744 | 2.98 | Upregulation |
| 10. | 5-enolpyruvylshikimate 3-phosphate synthase (ESPS) | 11286 | 13874 | 74066 | 2.17 | Upregulation |

### Transcription factors involved in sandalwood oil biosynthesis

Transcription factors are important regulators, which can regulate the development, maturation, oil biosynthesis and accumulation in plants [34,35]. Transcription factor database revealed 47 families of transcription factors in *Sa*SHc and 41 in *Sa*SLc distributed across the RNA-Sequence in sandalwood. Some of the abundant transcription factors included CDK7, ERCC2, ERCC3, CCNH, TAF8, TAF4, TFIIA, TFIIB, GTF2A, GTF2 and TBP (Table 7). Total fourteen upregulated transcription factors were identified *viz*, (1) transcription initiation factors TFIID subunit6, five folds in *Sa*SHc and four folds in *Sa*SLc (K03131, 0.86) (2) transcription initiation factor TFIID TATA-box-binding protein (K03120, 0.64) (3) transcription initiation factor TFIIA small subunit (K03123 FC 0.50) (4) transcription initiation factor TFIIF subunit α two copy (K03138, 0.44) (5) transcription initiation factor TFIIH subunit2 (K03142, 0.44), (6) cyclin-dependent kinase7 three copy in *Sa*SLc and one copy in *Sa*SHc (K02202, 0.42) (7) cyclin H one copy in *Sa*SHc and two copy in *Sa*SLc (K06634, 0.42) (8) CDK-activating kinase assembly factor MAT1 two copy in *Sa*SLc and one copy present in *Sa*SHc sample (K10842, 0.42) (9) transcription initiation factor TFIID subunit11 (K03135, 0.31) (10) transcription initiation factor TFIIF *β* subunit (K03139, 0.34), (11) transcription initiation factor TFIID subunit2 (K03128, 0.35), (12) transcription initiation factor TFIIE subunit *α*, two copy in *Sa*SHc (K03136, 0.24), (13) transcription initiation factor TFIIE subunit *β* (K03137, 0.27) (14) transcription initiation factor TFIID subunit 9B (K03133, 0.18) (Table S5). Nine genes were downregulated with FC range from −578 to −0.63. It included (1) DNA excision repair protein ERCC-3, 2 copy (K10844, −0.75), (2) transcription initiation factor TFIID subunit1 (K03125, −0.57), (3) transcription initiation factor TFIID subunit4, two copy (K03129, −0.17), (4) transcription initiation factor TFIID subunit12 (K03126, −0.17), (5) Transcription initiation factor TFIIA large subunit three copy in in both the accessions (K03122 −0.10), (6) transcription initiation factor TFIIH subunit 4 copy in *Sa*SHc (K03144, −1.0), (7) transcription initiation factor TFIIH subunit three copy in *Sa*SHc (K03143, −0.23), (8) transcription initiation factor TFIIB four copy in *Sa*SHc (K03124, −0.23), (9) transcription initiation factor TFIID subunit 5, two copy in *Sa*SHc and one copy present in *Sa*SHc sample (K03130 −0.63) (Table 7).

**Table 7.**
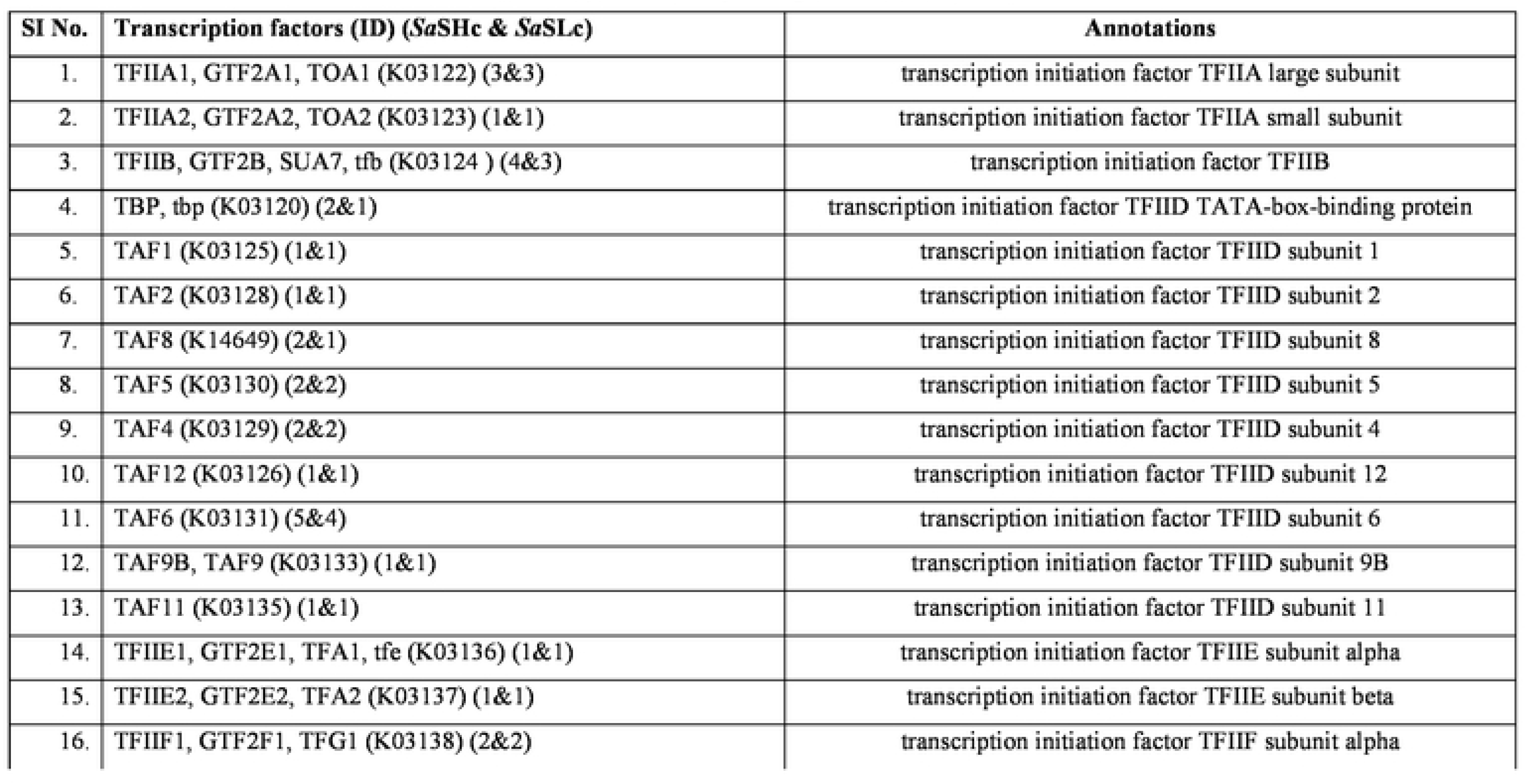
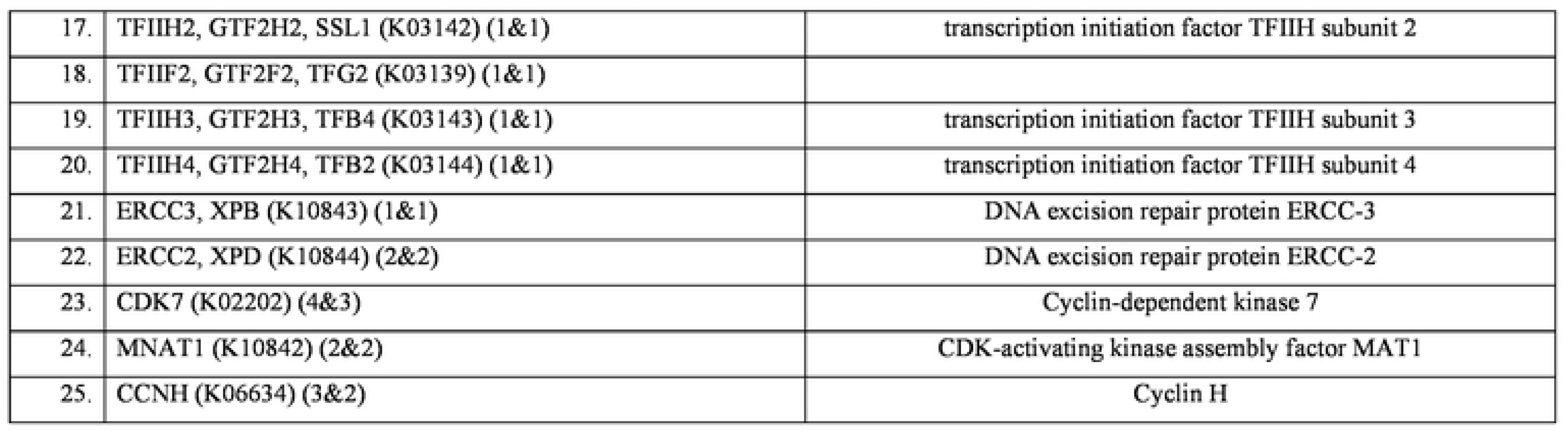
list of Transcription Factors (Ko3022) and genes encoding key enzymes for sandalwood oil biosynthesis whose expressions were altered in high oil (***Sa*SHc**) and low oil yielding (***Sa*SLc**) sandalwood

| SI No. | Transcription factors (ID) (*SaSHc* & *SaSLc*) | Annotations |
| --- | --- | --- |
| 1. | TFIIA1, GTF2A1, TOA1 (K03122) (3&3) | transcription initiation factor TFIIA large subunit |
| 2. | TFIIA2, GTF2A2, TOA2 (K03123) (1&1) | transcription initiation factor TFIIA small subunit |
| 3. | TFIIB, GTF2B, SUA7, tfb (K03124 ) (4&3) | transcription initiation factor TFIIB |
| 4. | TBP, tbp (K03120) (2&1) | transcription initiation factor TFIID TATA-box-binding protein |
| 5. | TAF1 (K03125) (1&1) | transcription initiation factor TFIID subunit 1 |
| 6. | TAF2 (K03128) (1&1) | transcription initiation factor TFIID subunit 2 |
| 7. | TAF8 (K14649) (2&1) | transcription initiation factor TFIID subunit 8 |
| 8. | TAF5 (K03130) (2&2) | transcription initiation factor TFIID subunit 5 |
| 9. | TAF4 (K03129) (2&2) | transcription initiation factor TFIID subunit 4 |
| 10. | TAF12 (K03126) (1&1) | transcription initiation factor TFIID subunit 12 |
| 11. | TAF6 (K03131) (5&4) | transcription initiation factor TFIID subunit 6 |
| 12. | TAF9B, TAF9 (K03133) (1&1) | transcription initiation factor TFIID subunit 9B |
| 13. | TAF11 (K03135) (1&1) | transcription initiation factor TFIID subunit 11 |
| 14. | TFIIE1, GTF2E1, TFA1, tfe (K03136) (1&1) | transcription initiation factor TFIIE subunit alpha |
| 15. | TFIIE2, GTF2E2, TFA2 (K03137) (1&1) | transcription initiation factor TFIIE subunit beta |
| 16. | TFIIF1, GTF2F1, TFG1 (K03138) (2&2) | transcription initiation factor TFIIF subunit alpha |
| 17. | TFIIH2, GTF2H2, SSL1 (K03142) (1&1) | transcription initiation factor TFIIH subunit 2 |
| 18. | TFIIF2, GTF2F2, TFG2 (K03139) (1&1) |  |
| 19. | TFIIH3, GTF2H3, TFB4 (K03143) (1&1) | transcription initiation factor TFIIH subunit 3 |
| 20. | TFIIH4, GTF2H4, TFB2 (K03144) (1&1) | transcription initiation factor TFIIH subunit 4 |
| 21. | ERCC3, XPB (K10843) (1&1) | DNA excision repair protein ERCC-3 |
| 22. | ERCC2, XPD (K10844) (2&2) | DNA excision repair protein ERCC-2 |
| 23. | CDK7 (K02202) (4&3) | Cyclin-dependent kinase 7 |
| 24. | MNAT1 (K10842) (2&2) | CDK-activating kinase assembly factor MAT1 |
| 25. | CCNH (K06634) (3&2) | Cyclin H |

**Table 8.**
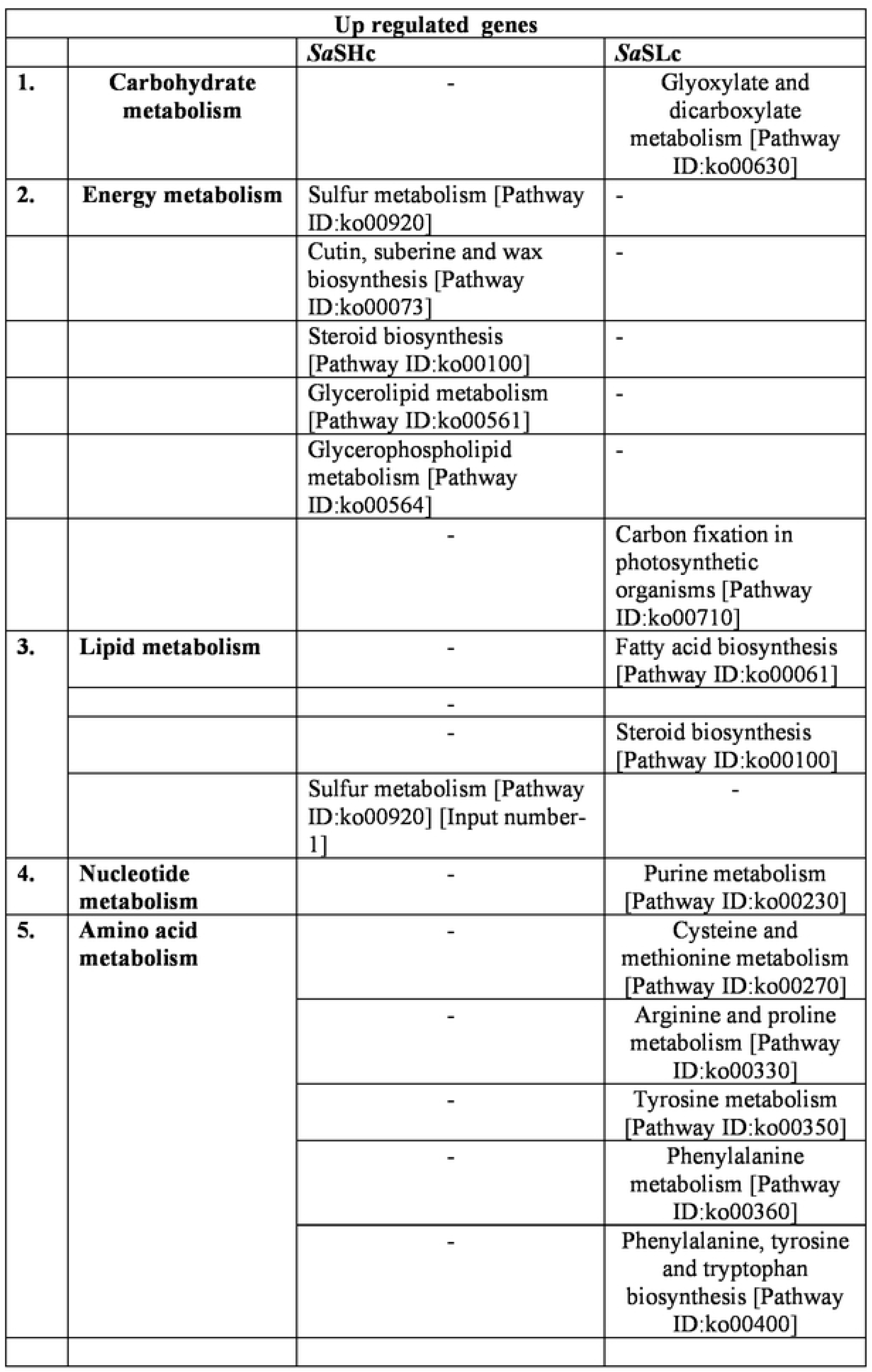
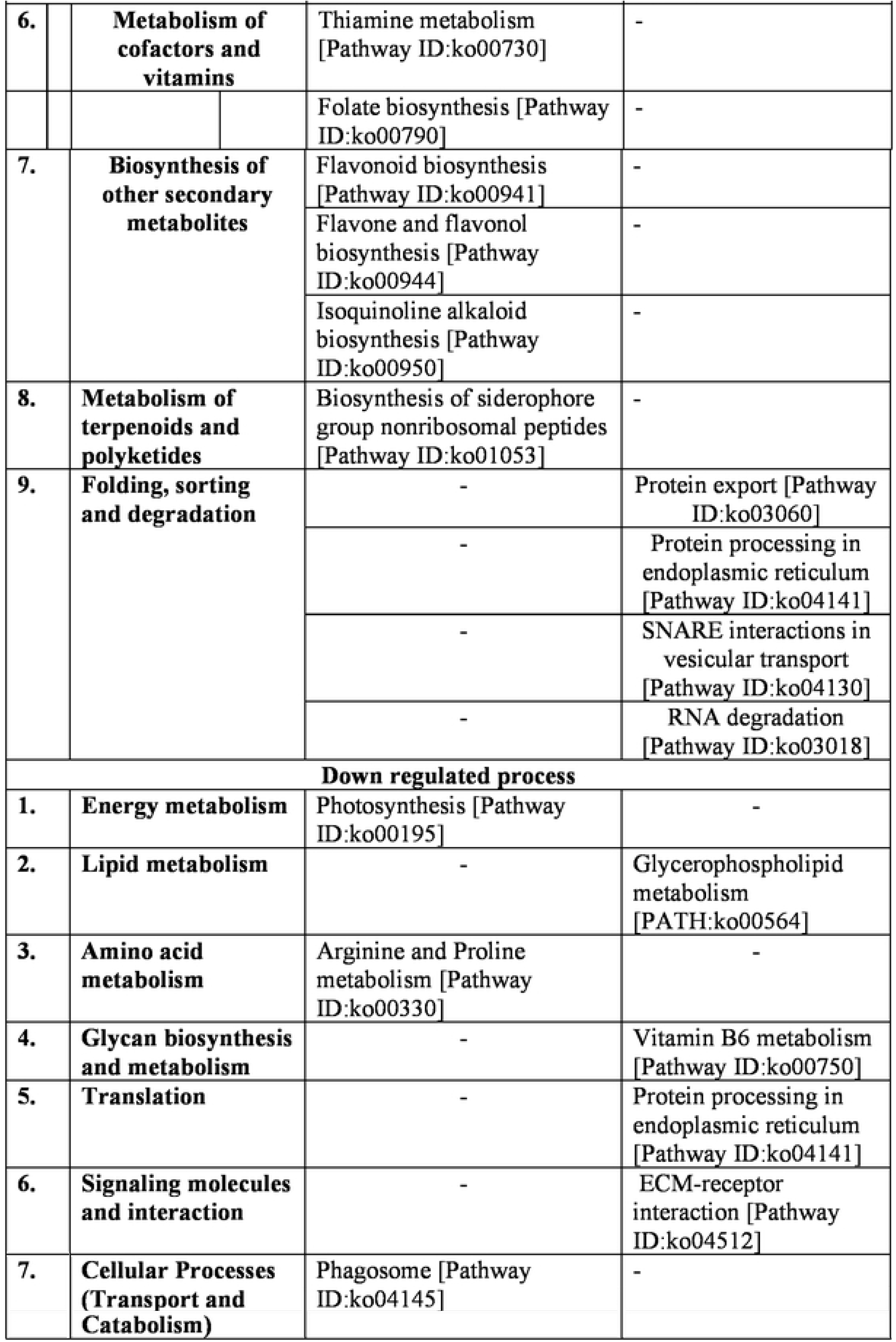
Total Biological process associated with differentially expresses genes (DEGs) in high and low oil yielding. *S. album* accessions

### Phylogenetic analysis of identified cytochrome family in RNA-seq of *S. album*

Cytochrome P450 mono-oxygenases putatively involved in sandalwood oil biosynthesis (Diaz-Chavez et al. 2013). In order to phylogenetic analysis of cytochromes, BLAST was performed on pooled RNA-seq data and total 237 cytochrome genes (FC 6.87-0.234) were listed in which 84 cytochrome genes were observed with FC>1.0. Based on their structures, total nine groups of cytochrome gens were resulted **i.** Cytochrome b561 **ii.** Cytochrome P450 **iii.** Cytochrome c oxidase **iv.** Cytochrome P45076C2 **v.** Cytochrome c oxidase subunit 1 **vi.** NADH-cytochromeb5 redutase **vii.** SaCYP736A167 **viii.** mitochondrial cytochrome b and **ix.** CytochromeP450 E-class (S6 Table).

### Distribution of shared gene clusters across plant species

In the current study, majority of the blast hits were found to be against *Vitis vinifera, Quercus suber, Juglans regia, Nelumbo nucifera, Thobroma cacao, Ziziphus jujuba, Hevea brasiliensis, Manihot esculenta* and *Jatropha curcus* (Fig 6). BLAST results were obtained for 91.77% of all the contigs with upregulated and downregulated genes (8.22% without BLAST hit). Whereby the 9 woody plant taxa *V. vinifera*: 4,710 (46.97%) *Q. suber*: 828 (8.25%), *J. regia*: 782 (7.82%), *N. nucifera*: 766 (7.64%), *T. cacao*: 460 (4.58%), *Z. jujuba*: 437 (4.35%), *H. brasiliensis*: 428 (4.26%), *M. esculenta*: 358 (3.57%), *J. curcus*: 338 (3.37%) and *A. thaliana* 23 (0.8%) with 896 genes were no blast hit were the species which gave the highest number of BLAST hits S6 Fig. Although many numbers of transcripts were not functionally annotated, this study provides more than 20,842 annotated transcripts, which can be directly used for further research in sandalwood species. Total 784 genes were upregulated and BLAST results were obtained for 770 (98.2%) genes were shared clusters with other plant species and 41 (5.2%) was found no blast hit S5 Fig. Total 339 genes were down regulated and BLAST results were obtained for 80.2% of all the contigs (19.2% without BLAST hit) S6 Fig.

**Figure 6.**
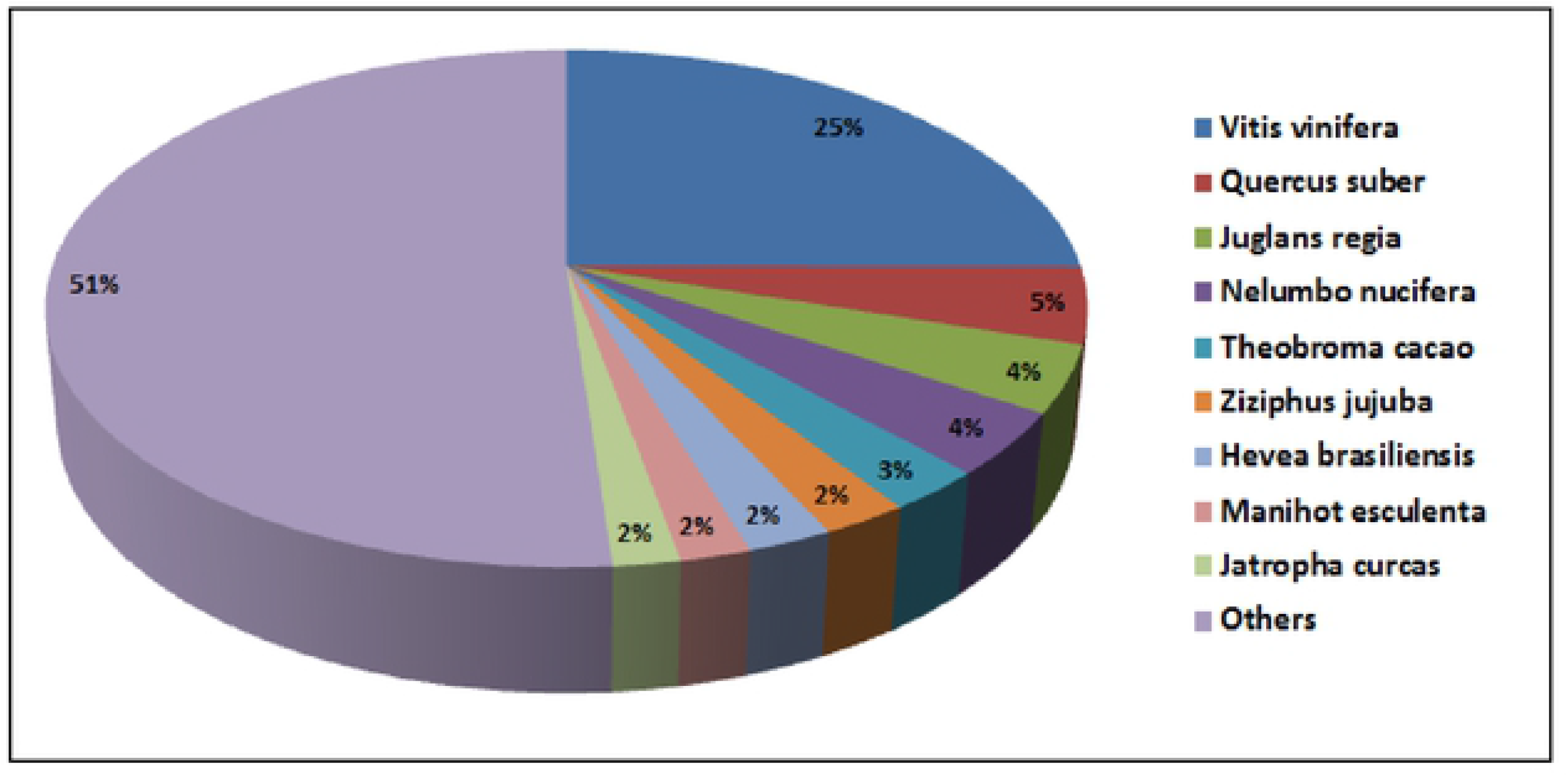
Top Blast Hit Species distribution of pooled CDS; Majority of the hits were found to be against *Vitis vinifera*.

### Validation of the expression profiles of candidate genes involves in high oil biosynthesis of sandalwood by Real Time PCR (q-PCR)

To validate the expression profiles of candidate genes obtained from the RNA-Seq analysis, six candidate genes relate with oil biosynthesis in the transition zone of sandalwood were selected for qRT-PCR analysis. The expression levels of the selected genes were compared with RNA-seq results. The expression patterns of RNA-Seq and qRT-PCR revealed that the expression pattern of these genes were consistent which indicated the reliability of the RNA-seq data Fig 7.

**Figure 7.**
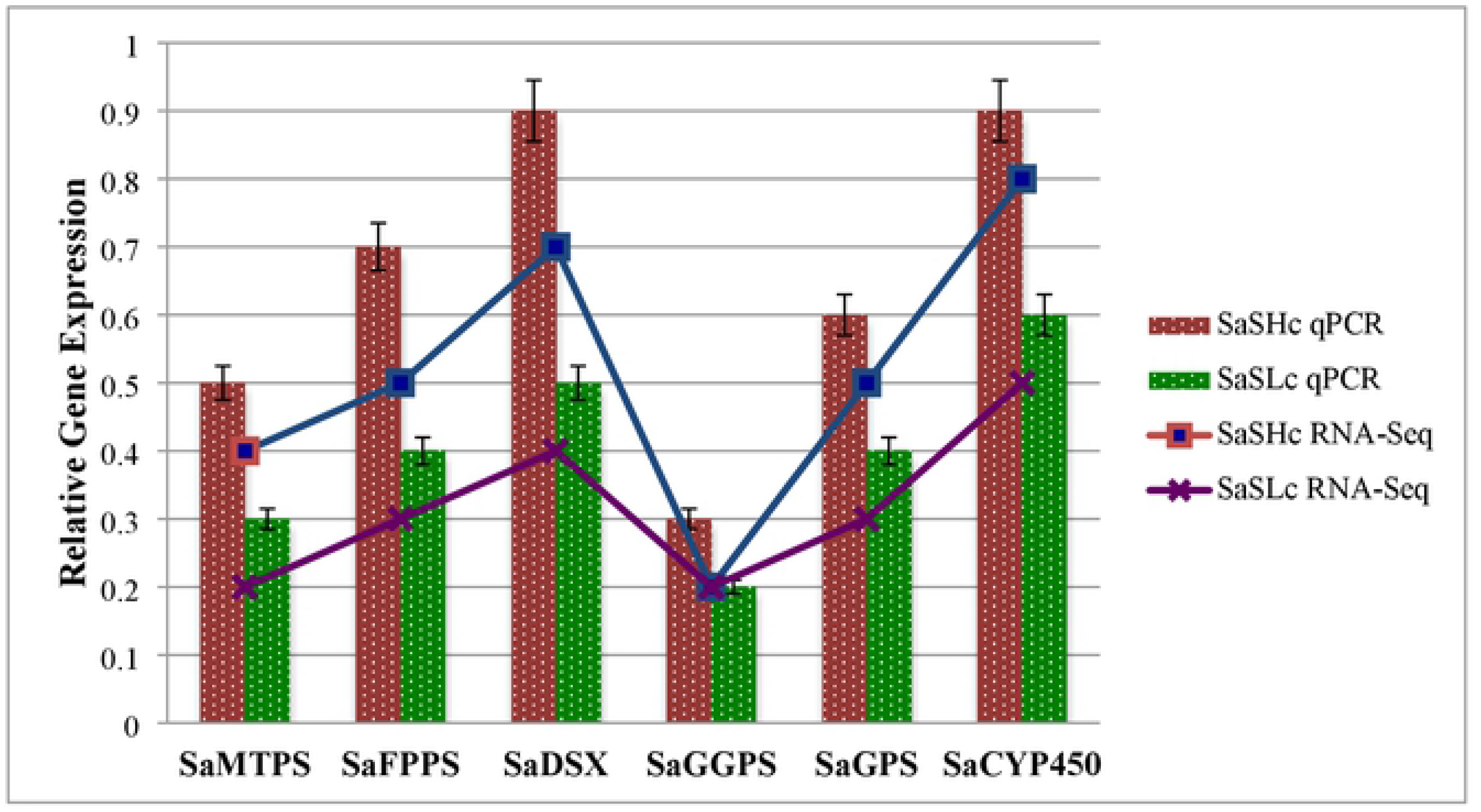
Validation of relative gene expression levels of DEGs by qRT-PCR. Purple and blue lines represents the RNA-Seq results, while red and green bars represent the qRT-PCR results. The error bars indicate the standard deviation.

## Discussion

*`Santalum album`* is a highly priced commodity and the tropical tree crop is facing a lot of problems in the country because of heavy occurrence of industrial uses and smuggling. It has been found that sandalwood oil of different accessions vary widely in terms of oil content with a negative correlation between heartwood and oil [8]. To understand the dynamic regulation of oil accumulation, comparative *`De novo`* transcriptome profiling of two identical accessions that differ significantly in oil content was carried out. Using comparative transcriptomics, we tried to infer the effect of change in gene structure difference in sandalwood accessions (*Sa*SHc and *Sa*SLc) Table 1; S1 Table S1. In recent years, RNA-seq has been extensively employed for sandalwood oil biosynthesis pathway [18,19,20]. Understanding the underlying molecular mechanism is important for developing high oil yielding cultivation of sandalwood. To the best of our knowledge, this is the first study reporting the comparative transcriptomic response of sandalwood using RNA-Seq approach and identified different group of genes in high oil yielding samples under the similar condition. The transcriptome assembly generated *Sa*SHc 3.95 billion and *Sa*SLc 2.89 billion raw reads with high PE reads transcripts, unigenes and CDS was observed Table 1; S3 Table. In other studies, on Santalum and other tree species similar results were observed like *S. album and Torreya grandis* [19,20,34,21,36]. Low raw reads were and low PE reads also observed in *S. album* [18,19,20,21]. Approximately 20,842 genes were annotated to a total of 22,710 GO terms annotations BLASTX hits against non-redundant plant species database (Table S3). Similar results were reported in in *Quercus pubescens* [37]. Approximately 65.29% genes were annotated in gene ontology terms and most of them were involved in process of cellular components followed by molecular functions Fig 1; Table 2. Further WEGO plot analysis showed that in cellular component *Sa*SHc had high number of genes than *Sa*SLc and the most enriched grouped in cell, cell part and membrane part. In molecular function of *Sa*SHc and *Sa*SLc most of the GO terms were involved in catalytic activity and binding. In biological processes majority of GO terms were grouped into two classes, metabolic and cellular process Fig 2 A & B. Similarly our data at GO level resemble previous work with morphophytes of vetiver, *Chrysopogon zizaniod*es [38].

The combined assembly of additional number of DEGs likely reflects the difference expression patterns between high oil yielding low oil yielding sandalwood accessions Fig 3. The combined assembly of sandalwood accessions revealed the change trends of DEGS in high oil biosynthesis is somewhat consistent with upregulation of candidate oil biosynthesis CYTP167 gene [18].

Various approaches for functional annotation of the assembled transcripts have been used to identify the genes in which mostly were involved in secondary metabolite biosynthesis in sandalwood. Overall 24 KEGG pathways were marked in this study, which were further categorized into five major pathways Table 3. To identify secondary metabolites and related metabolic pathways in respective samples DEGs were mapped to KEGG database and resulted 14 major pathways shown to play important role in sandalwood oil biosynthesis Table 5. Among them, high number of genes involved in terpenoid backbone biosynthesis followed by polyprenoid biosynthesis and carotenoid biosynthesis in *Sa*SHc accession Table 5. In contrast to the present study, low number of genes were involved in KEGG pathways in *S. album* [21,34].

In our study, relative gene expressions of sandalwood oil biosynthesizing genes listed in [13] were found upregulated (log2FC 1.0-3.5 Table 6.

We identified selected candidate genes which were specifically showed in *Sa*SHc were Cytochrome b4561, Geranyl-geranyl diphosphate synthase, Geranyl pyrophosphate synthase, Monoterpene synthase, Sesquiterpene synthase, Shikimate-O-hydroxy-cinnamoyl-transferase, E, E-farnesyl diphosphate synthase and De-oxy-D-xylulose-5-phosphate synthase (Table 6) along with previously identified genes [18,19,20,39]. The expression of *Sa*GGPS was found relatively high than other genes S4 Table. In *S. spicatum* two genes *viz,* santalene synthases and cytochromes P450 were reported [39]. In another study of *S. album*, low differential expression were observed in *Sa*DXS and *Sa*HMG-Co-A genes in callus whereas, expression level of *Sa*FPPS, *Sa*STPS and *Sa*MTPS were quantitatively found high in matured leaves of *S. album* [40] and high expression of *Sa*FDSE and *Sa*SS genes were reported in *S. album* transition zone of *S. album* [41]. The transcriptional mining identified number of transcripts, unigenes and CDS with log2 FC (0.67-3.55) GPS (6), GGPS (9), HMG-CoA (4), DXS (8), E-E-FDS (5), CYP450 (5), FPPS (10), PAL (4), MTPS (4) and ESPS (5) exhibited (Table 6; S4 Table). [42] Reported similar data in Chinese tree *Sindora glabra*. We identified several transcription families in our data set. But little is known about the transcriptional regulation of oil biosynthesis in sandalwood. Transcription factor database revealed 47 families of transcription factors in *Sa*SHc and 41 in *Sa*SLc distributed across the RNA-Sequence in sandalwood (Table 7; S5 Table). [21] Reported 58 families of transcription factors in RNA-Seq data of leaf of sandalwood. However, we were unable to detect some of the transcription factors in our data. The lower number presented in our data set is likely because we used core tissue of sandalwood for our transcriptome analysis. The oil biosynthesis genes were abundantly expressed in *Sa*SHc when compared to *Sa*SLc accessions and validated the participation of genes in high oil biosynthesis Table 5. We observed *Sa*CYP736A167 in our predicted gene sets, which identified as a candidate key oil biosynthesis gene in *S. album* in previous reports [18]. Phylogenetic analysis of RNA-Seq resulted nine groups of cytochromes in *Sa*SHc and six groups in *Sa*SLc S6Table. [21] identified 184 Cytochrome P450 in *S. album* genome and out of them, four genes were reported in [18,20]. The obtained result suggested that all cytochromes in *S. album* evolved from a common ancestor and closely related to each other. Overall 16,665 genes were found differentially expressed between *Sa*SHc and *Sa*SLc with high number of upregulated and low downregulated genes Fig. 3. Similar results were observed in *S. glabra*, *C. sinenesis, P. tomoentosa* [42,43,44]. However, low number of DEGs was reported by [20,34] in *S. album*.

Based on the functional annotation enrichment analysis of the differentially expressed genes, identified some overrepresented genes participated in high oil biosynthesis with the highest 96.46% similarity in cytochrome b560 and Cytochrome b561 containing protein At3g61750 with 67.43%. It is generally accepted that identification of orthologous gene clusters helps in taxonomic and phylogenetic classification. We identified, 11,013 orthologous gene clusters, suggested their conservation in the ancestry. The orthologous clusters of the transcriptome was observed among ten plant species *V. vinifera, Q. suber, J. regia, N. nucifera, T. cacao, Z. jujuba, H. brasiliensis, M. esculenta, curcus* and *A. thaliana* Fig 6; S4 Fig. Among them total 770 genes were found upregulated and 111 genes were downregulated S5 Fig. S6 Fig. However, [21] reported five plant species *viz, A. thaliana*, *C. clementine*, *P. Trichocarpa* and *V. vinifera* in *S. album*.

## Conclusion

The comparative analysis of the sandalwood oil accumulating core tissues of sandalwood showed that transcriptional regulation plays a key role in the considerable differences in oil content between high and low oil yielding sandalwood. The present study generated a well-annotated pair end read RNA libraries and the results unveiled genome wide expression profile of sandalwood oil biosynthesis. Analysis of transcriptome data sets, identified transcripts that encode various transcription factor, metabolism of terpenoids, environment response element and biosynthesis of other secondary metabolites. Nevertheless, we also discovered some of the oil biosynthesis candidate genes SaCYP736A167, DXR, DSX and FPPS genes that participates in sandalwood oil biosynthesis and accumulation of oil in heartwood. The results suggested an intricate signalling and regulation cascade governing sandalwood oil biosynthesis involving multiple metabolic pathways. These findings have improved our understanding of the high sandalwood oil biosynthesis at the molecular level laid a solid basis for further functional characterization of those candidate genes associated with high sandalwood oil biosynthesis in *S. album.* Understanding the molecular mechanism of high and low oil sandalwood by RNA-seq will lead to significant information for farmers and forest department. The accessibility of a RNA-Seq for high oil yielding sandalwood accessions will have broader associations for the conservation and selection of superior elite samples/populations for further multiplications.

## Acknowledgement

Authors are thankful to the Director, IWST, Group Co-ordinator Research, Head-Genetics and Tree Improvement Division, Institute of Wood Science and Technology and Data Computational Science Department Indian Institute of Science for encouragement to carryout the present study.

## Data archiving statement

The Transcriptome Sequence Read Archive (SRA) data of Sandalwood have been deposited in NCBI under Biosample accession: SAMN1569426 SRA accession number: PRJNA648820 (https://submit.ncbi.nlm.nih.gov/subs/bioproject/SUB7788726/overview).

## Author contributions

First author design the experiment, completed laboratory work and written manuscript. All other authors reviewed the manuscript and helped in formatting.

## Author statement

All authors read, reviewed, agreed and approved the final manuscript.

## Availability of data and materials

We declare that all data generated or analyzed during this study are included in this manuscript.

## Ethics approval and consent to participate

Not applicable.

## Consent for publication

Not applicable.

## References

1. Harbaugh DT, Baldwin BG. Phylogeny and biogeography of the sandalwoods (Santalum, Santalaceae); repeated dispersals throughout the pacific. Amer J of Bot. 2007; 94: 1028–1040.

2. Shashidhara G, Hema MV, Koshy B, Farooqi AA. Assessment of genetic diversity and identification of core collection in sandalwood germplasm using RAPDs. J Hort Sci Biotech. 2003; 78: 528–536.

3. Brand JE, Fox JED, Pronk G, Cornwell C. Comparison of oil concentration and oil quality from *Santalum spicatum*, *Santalum album* plantations, 8-25 years old, with those from mature *S. spicatum* natural stands. Australian Forestry, 2007; 70(4): 235−241.

4. Kumar ANA, Joshi G, Mohan Ram HY. Sandalwood: History, Uses, Present Status and the Future. Curr Sci. 2012; 103: 1408–416.

5. Moniodis J, Jones C, Renton M, Plummer J, Barbour E, Ghisalberti E. et al. Sesquiterpene Variation in West Australian Sandalwood (*Santalum spicatum*). Molecules. 2017; 22(12): 940.

6. Subasinghe SMCUP. Sandalwood Research: A Global Perspective. Journal of Tropi Fore and Envi. 2013; 3: 1–8.

7. Gowda, VSV. Global Emerging Trends on sustainable production of natural sandalwood. Proceedings of the Art and joy of wood conference, 19-22 October. Bangalore India. 2011.

8. Kumar ANA, Srinivasa YB, Joshi G, Seethram A. Variability in and relation between the tree growth, heartwood and oil content in sandalwood (*Santalum album* L.) Curr Sci. 2011; 100 (6):827–830.

9. Srimathi RA, Kulkarni HD. Preliminary finding on the heartwood formation in Sandal (*S. album* L.). Proceedings of the second forestry conference, Dehradun. Minor Forest Products II. 1980; 108–115.

10. Kulkarni HD, Srimathi RA. Variation in foliar characteristics in sandal. In Biometric Analysis in Tree Improvement of Forest Biomass (ed. Khosla, P. K.), International Book Distributors, Dehra Dun. 1982; 63–69.

11. Page T, Southwell I, Russel M, Tate H, Tungan J, Sam C, et al. Geographic and Phenotypic variation in heartwood and essential oil characters in natural populations of *Santalum austrocaledonicum* in Vanuatu. Chem Biodiv. 2010; 7:1990–2006.

12. Brand JE, Pronk GM. Influence of age on sandalwood (*Santalum spicatum*) oil content within different wood grades from five plantations in Western Australia. Aus Forest. 2011; 74:141–148.

13. Fatima T, Srivastava A, Somashekar PV, Vageeshbabu HS, Rao SM, Bisht SS (2019) Assessment of morphological and genetic variability through genic microsatellite markers for essential oil in Sandalwood (*Santalum album* L.). 3Biotech 9: 252.

14. Lee DJ, Burridge, AJ, Page T, Huth JR, Thompson N. Domestication of northern sandalwood (*Santalum lanceolatum*, Santalaceae) for indigenous forestry on the Cape York Penninsular. Aus Forest. 2018; 82 (S1): 14–22.

15. Rai, S. N., and Sharma, C. R. 1990. Depleting sandalwood production and rising prices. Indi Forest, 116, 348–355.

16. Zhang Y, Yan H, Li Y, Xiong Y, Niu M, Zhang, X, et al. Molecular Cloning and Functional Analysis of 1-Deoxy-D-Xylulose 5-Phosphate Reductoisomerase from *Santalum album*. Genes. 2021; 12, 626.

17. Jones CG, Keeling CI, Ghisalberti EL., Barbour, EL., Plummer, JA., Bohlmann, J. Isolation of cDNAs and functional characterisation of two multi-product terpene synthase enzymes from sandalwood, *Santalum album* L. Arch Biochem Biophys. 2008; 477:121–130.

18. Diaz-Chavez ML, Moniodis J, Madilao LL, Jancsik S, Keeling CI, Barbour EL, et al. Biosynthesis of Sandalwood oil: *Santalum album* CYP76F Cytochrome P450 Produce Santalols and Bergamotol. PloS One. 2013; 8: E75053.

19. Srivastava PL, Daramwar PP, Krithika R, Pandreka A, Shankar SS, Thulasiram HV. Functional characterization of Novel Sesquiterpene Synthases from Indian Sandalwood, *Santalum album*. Sci Rep. 2015; 5:10095.

20. Celedon JM, Chiang A, Yuen MMS, Diaz-Chavez ML, Madilao LL, Finnegan PM, Barbour EL, Bohlmann J. Heartwood specific Transcriptome and metaboloite signatures of tropical sandalwood (*Santalum album*) reveal the final step of (Z)-santalol fragrance biosynthesis. Plant J. 2016; 86: 289–299.

21. Mahesh HB, Subba P, Advani J, Shirke MD, Loganthan RM, Chandana S, et al. Multi-omics driven assembly and annotation of the sandalwood (*Santalum album*) genome. Plant Physio. 2018;176: 2772–2788.

22. Lardizabal K, Effertz R, Levering C, Mai J, M.C. Pedroso, Jury, T, Aasen E, Gruys K, Bennett K. Expression of *Umbelopsis ramanniana* DGAT2A in seed increases oil inSoybean. Plant Physio. 2008; 148, 89–96.

23. Shahid M, Cai G, Zu F, Zhao Q, Qasim MU. Hong Y., et al. Comparative Transcriptome Analysis of Developing Seeds and SiliqueWall Reveals Dynamic Transcription Networks for Effective Oil Production in *Brassica napus* L. Int J of Mol Sci. 2019; 20 (8):1982.

24. Rubio-Piña JA, Zapata-Pérez O. Isolation of total RNA from tissues rich in polyphenols and polysaccharides of mangrove plants. E J Biotech. 2011; 14: 5.

25. Fatima T, Srivastava A, Vageeshbabu S. Hanur VS, M. Rao MS. An Efficient Method to Yield High-Quality total RNA from wood tissue of Indian Sandalwood (*Santalum album* L.) suited for RNA-Seq Analysis. Ind Fores. 2021. In press.

26. Bolger AM, Lohse M, Usadel B. Trimmomatic: a flexible trimmer for illumina sequence data. Bioinfo. 2014;30: 2114–2120.

27. Henschel R, Lieber M, Wu L, Nista PM, Haas BJ, Leduc RD. Trinity RNA-Seq assembler performance optimization. XSEDE ’12: Proceedings of the 1st Conference of the Extreme Science and Engineering Discovery Environment: Bridging from the extreme to the campus and beyond July 2012. 2012; 45 : 1–8.

28. Li W, Godzik A. CD-hit: a fast program for clustering and comparing large sets of protein or nucleotide sequences. Bioinfo. 2006; 22(13):1658–1659.

29. Conesa A, Gotzs S, Garcia-Gomez JM, Terol J, Talon M, Robles M. Blast2GO: A universal tool for annotation, visualization and analysis in functional genomics research. Bioinfo. 2005; 21: 3674–3676.

30. Buchfink B, Xie, C, Huson, DH. Fast and sensitive protein alignment using DIAMOND. Nat Methods. 2015; 12(1):59–60.

31. Moriya Y, Itoh M, Okuda S, Yoshizawa AC, Konehia M. KAAS: an automatic genome annotation and pathway reconstruction server. W182– W185 Nucl Acids Res. 2007; 35: 182–185.

32. Anders S, Huber W. Differential expression of RNA-Seq data at the gene level–the DESeq package. Heidelberg, Germany: European Molecular Biology Laboratory (EMBL). 2012.

33. Howe EA, Sinha R, Schlauch D, Quackenbush J. RNA-Seq analysis in MeV, Bioinfo. 2010; 27(22): 3209–3210.

34. Li D, Jin C, Duan S, Zhu Y, Qi S, Liu K. et al. MYB89 Transcription Factor Represses Seed Oil Accumulation. Plant Physio. 2017; 173(2): 1211–1225.

35. Manan S, Chen B, She G, Wan X, Zhao J. Transport and transcriptional regulation of oil production in plants. Crit Rev in Biotech. 2017; 37(5): 641–655.

36. Zeng J, Chen J, Kou Y, Wang Y. Application of EST-SSR markers developed from the transcriptome of *Torreya grandis* (Taxaceae), a threatened nut-yielding conifer tree. PeerJ. 2018; 6: e5606.

37. Torre S, Tattini M, Brunetti C, Fineschi S, Fini A, Ferrini F, et al. RNA-Seq analysis of *Quercus pibescens* leaves: *De Novo* transcriptome assembly annotation and functional marker development. Plos One. 2014; 9: e112487.

38. Chakrabarty D, Chauhan PS, Chauhan AS, Indoliya Y, Lavania UC, Nautiyal CS. *De novo* assembly and characterization of root transcriptome in two distinct morphophytes of vetiver, *Chrysopogon zizaniod*es (L.) Roberty. Sci Rep. 2015; 5: 18630.

39. Moniodis J, Jones CG, Barbour EL, Plummer JA, Ghisalberti EL, Bohlmann J. The transcriptome of sesquiterpenoid biosynthesis in heartwood xylem of Western Australian sandalwood (*Santalum spicatum*). Phytochem. 2015; 113:79–86.

40. Misra BB, Dey S. 2013. Developmental variations in sesquiterpenoid biosynthesis in East Indian sandalwood (*Santalum album* L). Trees, 27: 1071–1086.

41. Rani A, Ravikumar P, Reddy MD, Kush A. Molecular regulation of santalol Biosynthesis in *Santalum album* L. Gene. 2013; 527: 642–648.

42. Yu N, Yang JC, Yin GT, Li RS, Zou WT. Transcriptome analysis of Oleoresin-Producing Tree *Sindora Glabra* and characterization of sesquiterpene synthases. Front of Plant Sci. 2018; 9: 1619.

43. Cao D, Liu Y, Ma L, Jin X, Guo G, Tan R, Liu Z, Zheng L, Ye F, Liu W Transcriptome analysis of differentially expressed genes involved in selenium accumulation in tea plant (*Camellia sinensis*). PLoS One. 2018; 13: e0197506.

44. Chen Z, Rao P, Yang X, Su X, Zhao T, Gao K, Yang X, An X. A Global View of Transcriptome Dynamics During Male Floral Bud Development in *Populus tomentosa*. Sci Rep. 2018; 8: 722.

